# *In vivo and in silico* experiment on *Mitragyna speciosa* offers new insight of antidiabetic potentials

**DOI:** 10.1101/2023.11.08.566341

**Authors:** Mohammad Uzzal Hossain, Md. Shahadat Hossain, Shajib Dey, A.B.Z. Naimur Rahman, Zeshan Mahmud Chowdhury, Arittra Bhattacharjee, Ishtiaque Ahammad, Md. Nazrul Islam, Md. Moniruzzaman, Md. Billal Hosen, Istiak Ahmed, Keshob Chandra Das, Chaman Ara Keya, Md. Salimullah

## Abstract

Diabetes Mellitus (DM) is a serious metabolic disease with several treatments available for managing it, however, they can be expensive and have side effects. Medicinal plants have been used for many years to treat numerous diseases. *Mitragyna speciosa (M. speciosa*) plant has been shown to have anti-diabetic properties in preclinical studies. Hence, this study aimed to investigate the phytochemical compositions and anti-diabetic effects of *M. speciose,* using both *in vivo* and *in silico* approaches. In the in vivo study, experimental diabetes was induced in Swiss albino mice using the drug alloxan (150 mg/kg). Four groups of diabetic mice were taken. Two of the groups were given extracts at doses of 200 mg/kg and 400 mg/kg respectively. Diabetic mice treated with the reference drug, glibenclamide (5mg/kg) were chosen as a positive control, and mice with only vehicles were considered as a negative control. The Oral Glucose Tolerance Test (OGTT) and the acute toxicity test were performed. In the *in silico* study, molecular docking and dynamics were performed for the identification of the plant compounds that could effectively bind with the DPP4 receptor. Analysis of the study suggested that the lethal dose (LD50) values were greater than 2000 mg/kg, indicating that a dose below this level can be selected. The OGTT results showed that both doses of *M. speciose* extracts significantly reduced blood glucose levels (P<0.0001). However, neither dose exhibited a significantly higher blood glucose reduction compared to glibenclamide (5 mg/kg) (P > 0.05). Phytochemical screening and the ADMET profile analysis suggested four key compounds: Mitragynine, Corynantheidine, Corynoxine, and Speciociliatine in *M. speciosa.* The molecular docking analysis revealed these four compounds as potential antidiabetic agents considering their high binding affinity to the DPP4 receptor. These compounds were also found to be stable in the DPP4 binding pocket, as evidenced by the molecular dynamics simulation. Lastly, it can be demonstrated that the *in silico* experiments confirmed *M. speciosa* extracts to be capable of reducing blood glucose levels because of the presence of compounds.

## Introduction

Diabetes mellitus (DM) is a set of metabolic disorders where the blood glucose levels remain elevated or hyperglycemic for an extended duration. The most widespread type of diabetes is Type 2 diabetes mellitus (T2DM), which accounts for 95% of all cases in contrast to Type 1 diabetes (T1DM) [1]. Type 1 diabetes develops when the body is unable to produce insulin, while in type 2 diabetes, the body becomes resistant to insulin. Additionally, changes in multiple genes and their products can potentially contribute to the development of type 2 diabetes [2], [3]. Effective management of type 2 diabetes typically involves improving insulin sensitivity and reducing hyperglycemia through lifestyle modifications, pharmacological treatments, and monitoring of blood glucose levels [4]. Several recent studies have highlighted the importance of insulin therapy for the effective management of diabetes [5]. However, there are still some challenges associated with insulin therapy, including the potential for weight gain, hypoglycemia, and injection site reactions. Over the past few decades, DM treatment methods have been substantially improved [6]–[8].

Anti-diabetic medications, however, can have substantial side effects, including hypoglycemic coma, and liver and kidney diseases [9]. According to the World Health Organization (WHO), adding medicinal plants to the diet can help manage diabetes [10], [11]. A total of four billion people in developing countries use medicinal plants to treat diseases such as DM[12], [13]. Preclinical and clinical studies have demonstrated the effectiveness of herbs, vitamins, and plant compounds in lowering blood sugar [14], [15]. For example, researchers have found that zinc intake controls insulin receptors and elevates insulin activity [16]. Another study showed that garlic prevents diabetic retinopathy in adult albino rats [17]. The antidiabetic phytochemicals have been identified based on the differences in their chemical structures [18], [19]. Some of the major classes of phytochemicals are Alkaloids, aromatic acids, carotenoids, coumarins, essential oils, flavonoids, glycosides, organic acids, phenols and phenolics, phytosterols, protease inhibitors, saponins, steroids, tannins, terpenes, and terpenoids [20]–[24].

Despite the promising findings, plant-derived compounds with pharmacoinformatics elucidation for the identification of novel therapeutics against diabetes have been understudied, and further research is needed to fully understand the therapeutic potential and safety of *M. speciosa* [30], [31]. Previous studies have been confined to the identification of the compounds of *Mitragyna speciosa,* their pharmacological effects, and their clinical implications [32]. However, the present study will provide the opportunity to explore new potential compounds from *M. speciosa* and to reveal the molecular mechanism of these compounds in the receptor.

*M. speciosa* and the other species of the genus *Mitragyna* share several common phytochemical compounds [33]. The use of Mitragyna species in the treatment of diabetes has been supported by a number of ethnobotanical *in vitro and in vivo* evidence, although this claim has not been scientifically confirmed [34]. Therefore, the purpose of this study was to ascertain the scientific basis for the traditional use of *M. speciosa* in the treatment of diabetes using methanolic extracts on diabetic mice that had received an alloxan injection. We then revealed the mechanism of action of the compounds of *M. speciosa* in the receptors in terms of their binding affinity and stability through computational studies. This comprehensive study will provide insights into the enhanced perception of the drug-likeness of the phytochemicals from kratom and the drug-protein interaction of these compounds for the future blueprint to be contemplated as antidiabetic drugs.

## 2. Methods

### 2.1 Collection, Preparation, and extraction of plant material

The *M. speciosa* leaves were collected from Rangamati, Bangladesh. The Plant Biotechnology Division of the National Institute of Biotechnology performed the taxonomic identification. The leaves of *M. speciosa* were set to air dry at room temperature in a shady spot for a week. For the extraction process, the dried plant material was manually ground into a fine powder. The dried and powdered *M. speciosa* leaves were properly dissolved in methanol and then left at room temperature for 72 hours with constant magnetic stirring. The mixture was then extracted from the methanolic extract and filtered. About 100g of the dried methanolic crude extract was collected. For the experiment, the freeze-dried methanolic extract was preserved in the refrigerator. After that, the crude extract of dried plants was reconstituted in a suitable solvent for oral use.

### 2.2 Experimental animals

We used adult Swiss albino mice that were eight weeks old and weighed between 24 and 35 g. The animals were raised in the animal house using polypropylene cages at a temperature of 23 ± 2°C with a 12 h/12 h light/dark cycle [35]–[38]. The animals were given regular food pellets and water throughout the experiment, with the exception of the fasting period. The study protocol was approved by the Plant Biotechnology Division of the National Institute of Biotechnology, and the animals were handled and cared for in compliance with the ethical guidelines for the use of lab animals.

### 2.3 Induction of experimental diabetes

A glucometer was used to assess the weight and fasting blood glucose levels of Swiss albino mice after they had been fasting for 12–14 hours. Following this, a single intraperitoneal injection of fresh alloxan monohydrate solution (150 mg/kg body weight) was administered to the mice, leading to the development of diabetes. Alloxan was prepared with 0.5 mL of sodium citrate at pH 4.5 prior to injection. The animals were then supplied with food and water after thirty minutes of alloxan induction [39], [40]. Following a 48-hour alloxan injection, the plasma blood glucose levels of each animal were measured using tail blood, and the blood glucose levels of animals with fasting were greater than 200 mg/dL [41], [42].

### 2.4 Oral glucose tolerance test (OGTT)

The animals were kept fasting but had access to water for 24 hours before conducting the OGTT experiments. A 1 mL/kg dose of oral glucose solution (2 g/kg body weight) was given. Blood glucose levels were obtained at 0, 30, 60, 90, and 120 minutes after the glucose was administered [35].

### 2.5 Acute toxicity test

Acute toxicity tests on the crude extract were performed after the animals had fasted the previous night and only consumed water [43]. Five groups were created, and each group had eight mice. The vehicle was provided only to the control group. 1000, 2000, and 4000 mg/kg of crude extract of *M. speciosa* were given to the treatment group [44]. Animals were closely observed for any visible toxicities or unusual behaviors for the first four hours after receiving the extract, such as agitation, tremors, diarrhoea, lethargy, weight loss, or paralysis [45]. After that, they were observed every day for two weeks to check if there had been any changes to their general behavior or other physical activities. Food became available 4 hours after extract administration.

### 2.6 Experimental Design

The five groups of mice (both male and female) were selected for this study. Group 1 mice were non-diabetic and served as the naive control group, receiving only the vehicle DMSO. Group 2 mice were diabetic and were employed as a negative control along with a vehicle. Group 3 is a positive control including a vehicle and glibenclamide (5 mg/kgb.w). Group 4 received 400 mg/kgb.w of the extract, while Group 5 received 200 mg/kgb.w of the extract in a vehicle [42].

### 2.7 Virtual screening of phytochemicals

The plant extracts were subjected to preliminary phytochemical screening using standard protocols to determine the presence or absence of secondary metabolites such as alkaloids, steroidal compounds, phenolic compounds, flavonoids, saponins, tannins, and anthraquinones. The plant compounds of *M. speciosa* leaves that may serve as anti-diabetics, as well as the other compounds discovered through laboratory study, were identified through extensive literature research. Seven (7) compounds of *M. speciosa* leaves were selected after extensive investigation, while others were discarded. The two-dimensional (2D) and three-dimensional (3D) structures of the compounds were extracted from the PubChem database [46].

### 2.8 Preparation of Target Protein

The crystal structure (3D) “5T4H” of the target protein dipeptidyl peptidase 4 (DPP4) of *Homo sapiens* was retrieved in the protein data bank (PDB) of the Research Collaboratory for Structural Bioinformatics (RCSB) PDB [47]. The PDB structure was cleaned up using the Discovery Studio 4.0 client to get rid of the connected ligands, water atoms, hetams, and A chain (**Supplementary Figure 1**).

### 2.9 Pharmacoinformatics Elucidation (ADMET Profiling)

The ADME (absorption, distribution, metabolism, and excretion) profiles of seven active compounds found in *M. speciosa* leaves were identified using ADMETlab 2.0 and the pkCSM online servers [48], [49]. The distribution features of these compounds were examined using pkCSM, while the adsorption, metabolism, and excretion of diverse substances were investigated using ADMETlab 2.0. Graph-based signatures were used by the pkCSM to identify ADMET characteristics and support drug development. The ProTox-II server was used to calculate seven toxicity parameters, including hepatotoxicity, carcinogenicity, immunotoxicity, and mutagenicity [50]. The ProTox-II service uses a variety of machine learning methods, molecular similarity, and pharmacophore analysis to predict various toxicity endpoints.

### 2.10 Energy minimization of the protein

Using a variety of force folds, the Swiss PDB Viewer was used to minimize the energy of the chosen 3D structures [51]. The software allowed for the minimization and computation of the electrostatic and van der Waals energy of the protein. With the help of van der Waals and electrostatic energies, the protein was able to achieve a stable, low-energy state. The potential energy of the protein was also examined using the reduction approach for better accuracy of the protein structure and function.

### 2.11 Structural optimization of the compounds

The Avogadro program was utilized to optimize the selected compounds, providing the compounds with better arrangements with the lowest energy [52]. The optimized structures serve as a starting point for further computations, including docking investigations, quantum chemistry calculations, and simulations of molecular dynamics. By using Avogadro for structural optimization, we were able to produce novel compounds with the needed attributes while also guaranteeing the dependability and accuracy of our findings.

### 2.12 Active Site Analysis

CASTp was utilized to locate the active site and pocket of the DPP4 protein [53]. This program employs a sophisticated algorithm to calculate the three-dimensional solvent-accessible surface area of the protein, giving users a comprehensive view of the active site, including the precise location of the pocket and potential binding sites for small molecules.

### 2.13 Molecular Docking Analysis

Molecular docking is a computational technique for predicting the binding of small molecule ligands (such as drug candidates) to a receptor or protein target of interest [54]. In our research, we utilized AutoDock tools to simulate docking [55]. In the beginning, we added polar hydrogen to our target receptor protein DPP4 (PDB: 5T4H_B) and saved it as a PDBQT file in AutoDock. In order to observe its interaction with drug compounds, the prepared crystal structure of 5T4H_B was then covered with a grid box parameter based on the binding site residues. For each compound, the dimensions (X = 70, Y = 38, and Z = 50) and center (X = −11.527, 57.458, and Z = 27.219) of the grid box were set, as well as the grid box spacing (1.0). The torsional bonds were then liberated from the ligands and formatted as PDBQT files. pyMOL and Discovery Studio were used to visualize the protein-ligand interactions [56], [57].

### 2.14 Molecular Dynamics

DPP4 and ligand complexes stability (Mitragynine, Corynantheidine, Corynoxine, and Speciociliatine) stability was determined under specific physiological conditions. A 100 ns simulation was performed with GROningen MAchine for Chemical Simulations aka GROMACS (version 5.1.1) [58]. The simulation information was gathered to calculate the root mean square deviation (RMSD), root mean square fluctuation (RMSF), the radius of gyration (Rg), and solvent-accessible surface area (SASA) data. MD simulations were performed on the “bioinfo-server” running on the Ubuntu 18.4.5 operating system located at the Bioinformatics Division, National Institute of Biotechnology. Graphs were visualized and plotted using the QtGrace package visualization tool.

### 2.15 Statistical analysis

The GraphPad Prism software was used for the statistical analysis. An Analysis of Variance (ANOVA) was carried out between the groups, and a P-value of less than 0.05 was considered to be statistically significant [59].

## 3. Results

The overall workflow has been demonstrated in **Figure 1** and the location of the sample collection has been shown in **Figures 2A and 2B**.

**Figure 1:**
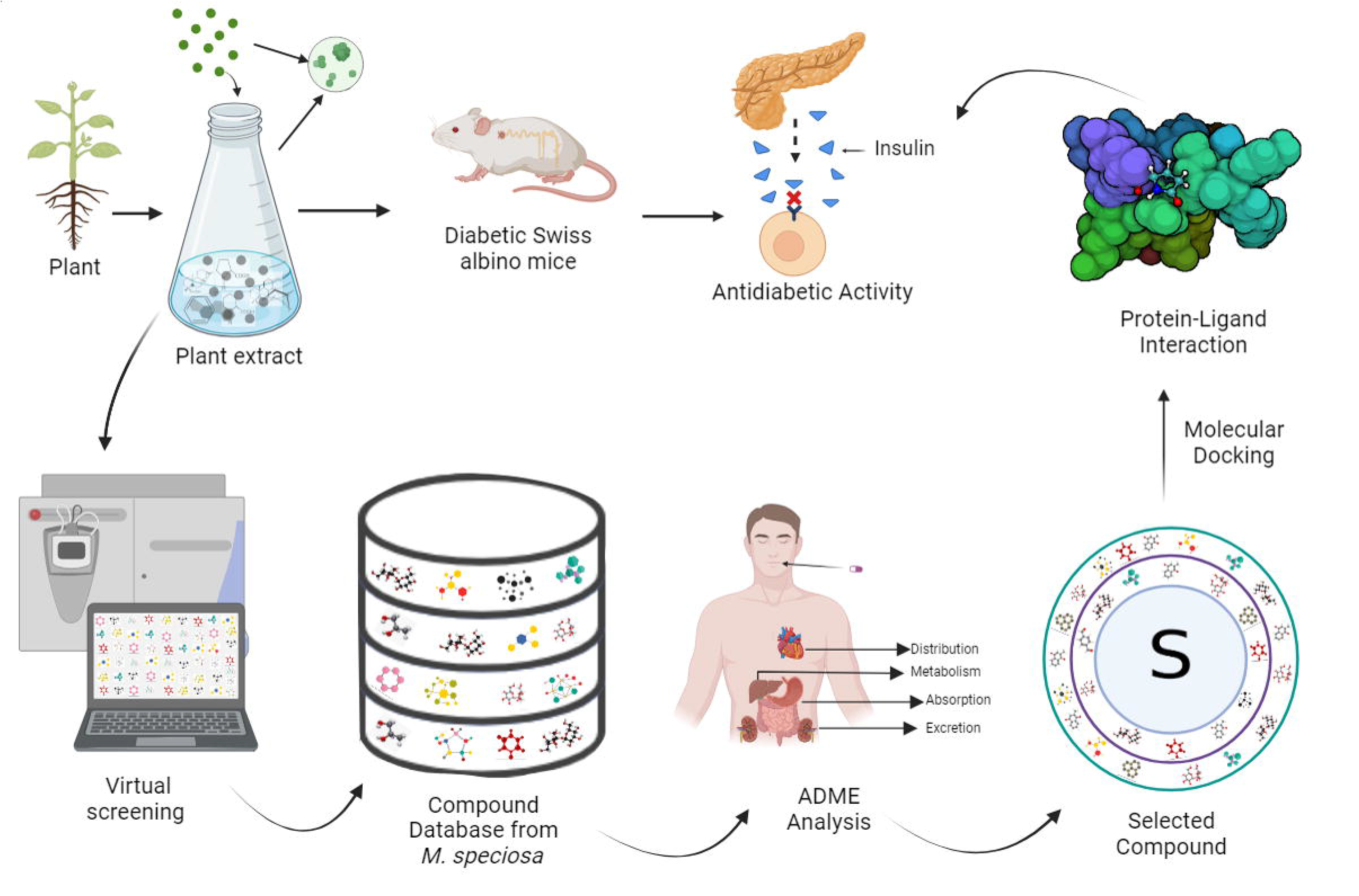
Schematic representation of the study

**Figure 2:**
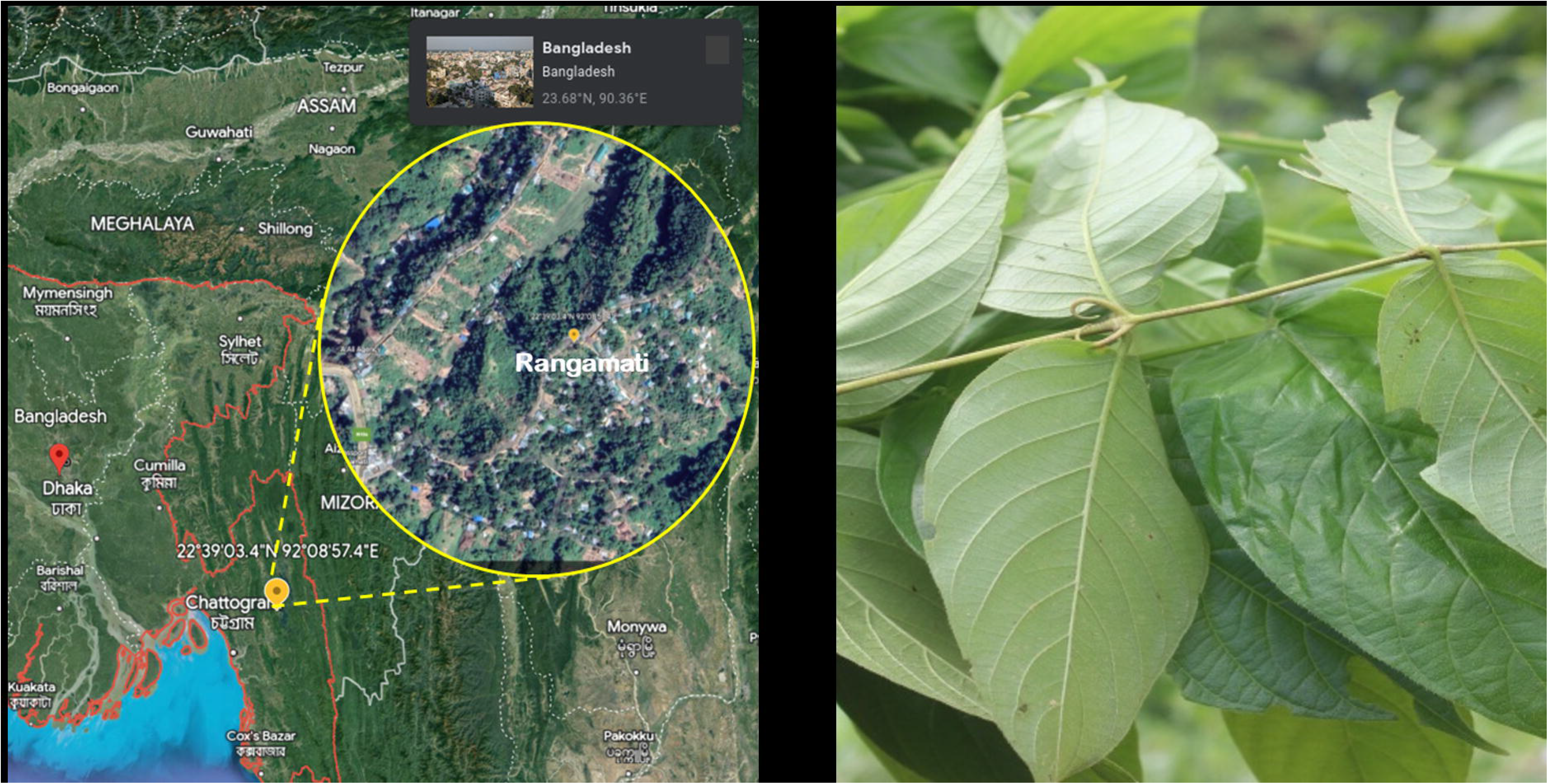
2A) Location of the sample collection. 2B) Plant parts (leaf and stem) pictorial view of the sample

### 3.1 OGTT and dose selection

For this investigation, male and female mice with blood glucose levels higher than 200 mg/dL were selected. The OGTT experiments yielded intriguing results in non-diabetic mice, as a significant difference was observed after the glucose induction in the response of mice at given intervals of 60, 90, and 120 minutes. However, the baseline blood glucose level (BGL) was reached in non-diabetic mice after 180 minutes, as expected **(Figure 3A and 3B).** In contrast, diabetic mice were not able to reach their baseline blood glucose levels at 180 minutes, despite the blood glucose levels being high at 60 minutes, 90 minutes, and even 180 minutes (**Figure 3A and 3B**). The area under the curve (AUC) of the control and treated mice was also significant (**Figure 3C and 3D**). Since the extracts did not manifest any explicit toxicity signs in mice, they were confirmed to be safe up to a dose level of 4000 mg/kg body weight. Two weeks later, the alterations were carefully observed, and it was found that there had been no significant changes in the animal’s sensory or motor ability, weight, sluggishness, respiration, paralysis, convulsions, or mortality.

**Figure 3:**
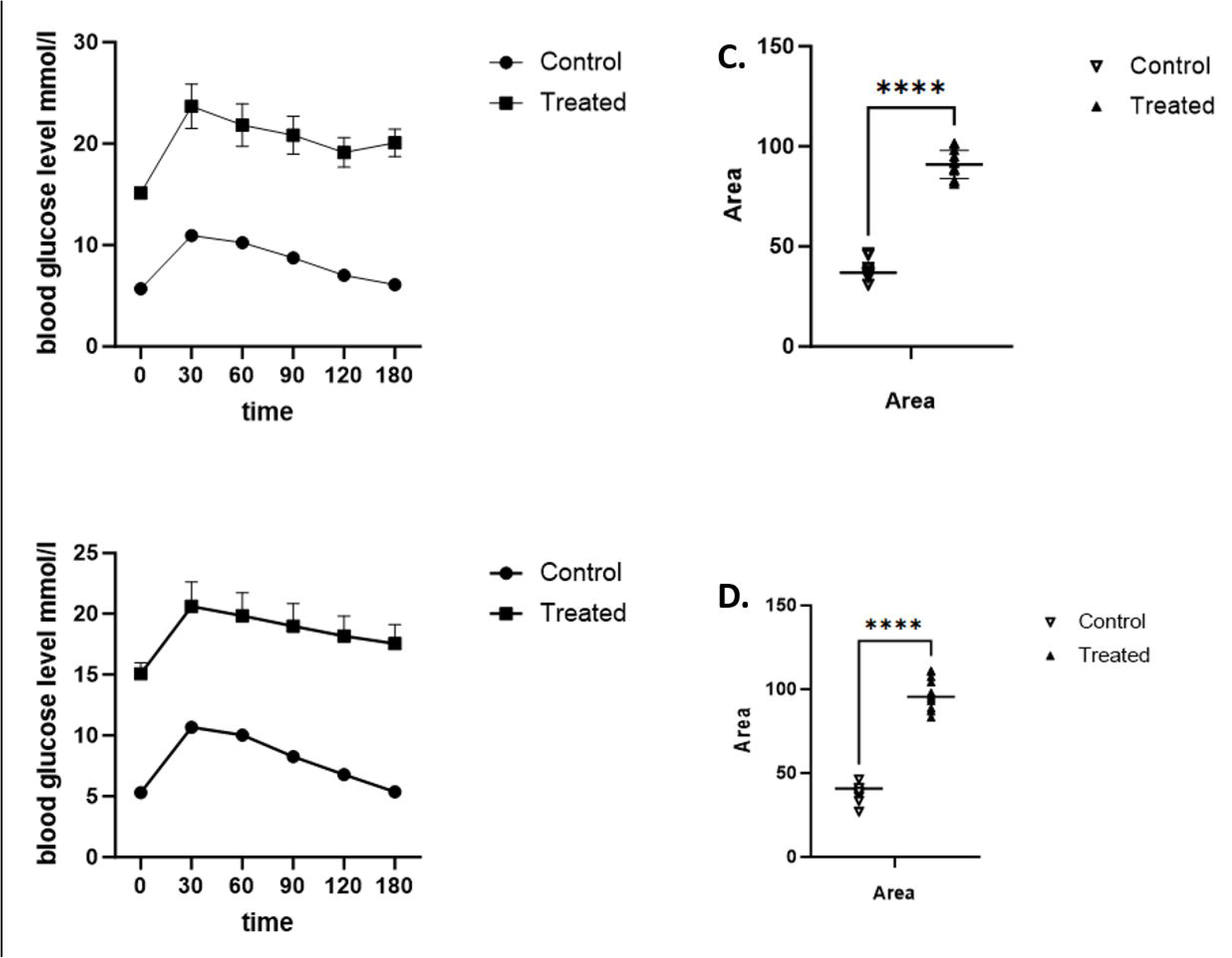
Oral glucose tolerance test (OGTT) of alloxan induced diabetic mice and non-diabetic mice. Mice (n=48) were fasted overnight. Glucose was administered at T=0 and at a level of 3g/kgb.w. Blood glucose levels were evaluated thirty minutes prior to dosing 30, 60, 90, 120, and 180 minutes. A) OGTT test in male mice showed high significance at each time point in multiple t-test analysis. The area under the curve (AUC) of each blood glucose level value was collected and found highly significant in control vs treated male mice (B). C) OGTT test in female mice and D) AUC in female mice (glucose treated) has also shown high significance in comparison to control mice. Data are mean ± SEM; **p<0.01, ****p<0.0001 compared to vehicle (N=6).

### 3.2 Effect of Plant Extracts on Blood Glucose Levels in Alloxan-induced Diabetes in Mice

The blood glucose levels were measured in 5 groups of male and female mice. The extracts treated groups (200mg/kg and 400mg/kg) significantly reduced BGL compared with the diabetic control group at day 7. According to the results, the BGL of the glibenclamide-treated group was also significantly lower on the seventh day (P > 0.001) than the diabetic control group. The results suggested that there were no significant differences between the glibenclamide-treated group and plant extracts-treated groups. However, 200mg/kg and 400mg/kg doses did not show any significant difference (**Figure 4A and 4B**).

**Figure 4:**
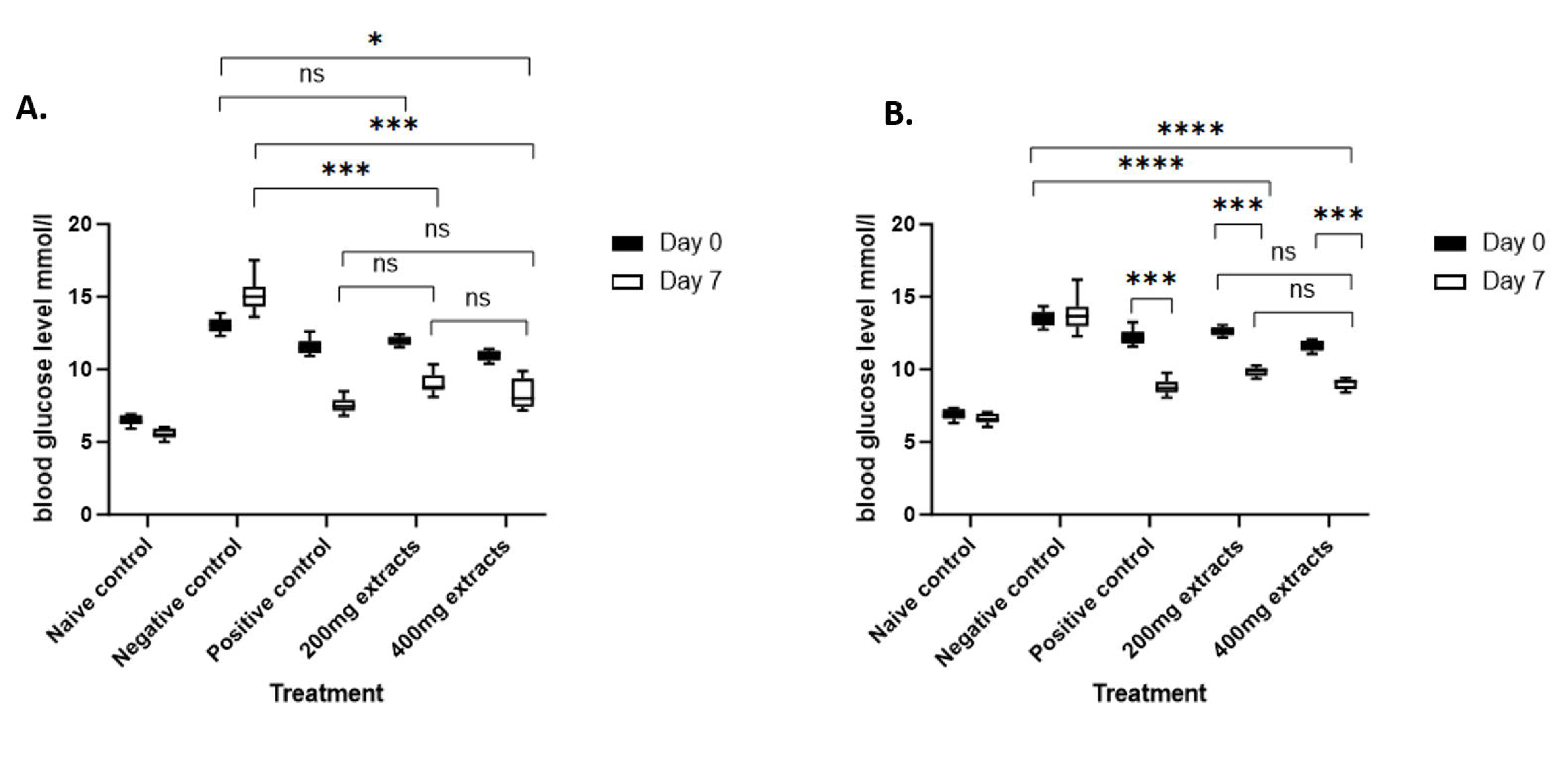
Effect of methanolic extract of *M. speciosa* leaves in both male (4A) and female mice (4B). Five (5) groups of mice consisting of 8 mice n=40 (male) and n=40 (female) were used for 5 types of experiments. Naïve control (nondiabetic with the vehicle), negative control (diabetic with the vehicle), positive control (5mg/kgb.w glibenclamide with the vehicle in alloxan-induced diabetic mice), and 200mg/kgb.w and 400mg/kgb.w of methanolic extracts were used in mice. The effect of methanolic extracts in male were analyzed in two ways (Effect vs Treatment) followed by Tukey’s multiple experiment. Data are presented as mean values ± SEM. **** p<0.0001; *** p<0.001; ** p<0.01; * p<0.05; ns= non-significant.

### 3.3 Retrieval of active compounds of *Mitragyna speciosa* leaves

A Literature study was conducted, and the list of bioactive molecules in the leaves of *M. speciosa* was found in the previous studies. The primary group of bioactive compounds in *M. speciosa* leaves are primarily alkaloids, flavonoids, terpenoids, and polyphenols (**Supplementary Table 1**).

### 3.4 ADME and Toxicity Analysis

The present study has selected seven compounds, such as Mitragynine, 7-Hydroxymitragynine, Corynoxine, Speciociliatine, Isorhyncophylline, Corynantheidine, and Isomitaphylline for pharmainformatics elucidation. The ADMET properties of seven compounds were analyzed to gain insight into their pharmacokinetic and toxicological properties (**Table 1**). Mitragynine and Corynantheidine, with a value of 0.005, exhibited the lowest intestinal absorption among all the selected compounds. However, 7-hydroxymitragynine had the highest Caco-2 permeability value with a value of −4.655, indicating a greater intestinal absorption potential. Corynoxine and Isorhyncophylline showed the highest values for blood-brain distribution at 0.01, while Corynantheidine had a moderate value of 0.442. Isorhyncophylline and Isomitraphylline posed the highest unbound fractions in plasma, with values of 0.353% and 0.357%, respectively. Corynantheidine had the highest volume of distribution (Vd) at 1.35 L/kg, while 7-Hydroxymitragynine had the lowest at 0.74 L/kg. All compounds had negative CNS permeability values, indicating that they could not easily cross the blood-brain barrier. In terms of metabolism, Corynoxine and Isomitraphylline possessed the lowest CYP1A2 inhibition potential, whereas 7-hydroxymitragynine and Isomitraphylline possessed the lowest CYP1A2 substrate potential. Isomitraphylline had the highest value for CYP2C9 inhibition at 0.356, while 7-Hydroxymitragynine had the lowest value at 0.131. Mitragynine and Speciociliatine had the highest CYP2C9 substrate potential at 0.658, while Corynoxine and Isorhyncophylline had the lowest at 0.220. Isomitraphylline exhibited the highest clearance (CL) rate (11.535), while Corynantheidine exhibited the lowest (4.934). Isomitraphylline had the shortest half-life at 0.243 hours, while Mitragynine and Spectrocilatine had the longest at 0.38 hours. In addition, all compounds have a high BBB (blood-brain barrier) score, indicating their potential to cross the blood-brain barrier and enter the central nervous system. In terms of distribution, all six compounds have a small unbound fraction in plasma, indicating high protein binding. Mitragynine and Corynoxine have a high volume of distribution (Vd) (L/kg), whereas 7-Hydroxymitragynine, Speciociliatine, Isorhyncophylline, and Isomitraphylline have lower values, indicating a more restricted distribution. In addition, each of the six substances has a low CNS permeability, indicating that they are unable to cross the blood-brain barrier. Some of these compounds are more susceptible than others to inhibition or induction of cytochrome P450 enzymes during metabolism. Mitragynine and Corynoxine, for example, inhibit CYP enzymes to a relatively low degree, whereas 7-Hydroxymitragynine, Speciociliatine, Isorhyncophylline, and Isomitraphylline have higher inhibition scores. All six compounds have relatively short half-lives in terms of excretion, with Isomitraphylline having the shortest half-life.

**Table 1:**
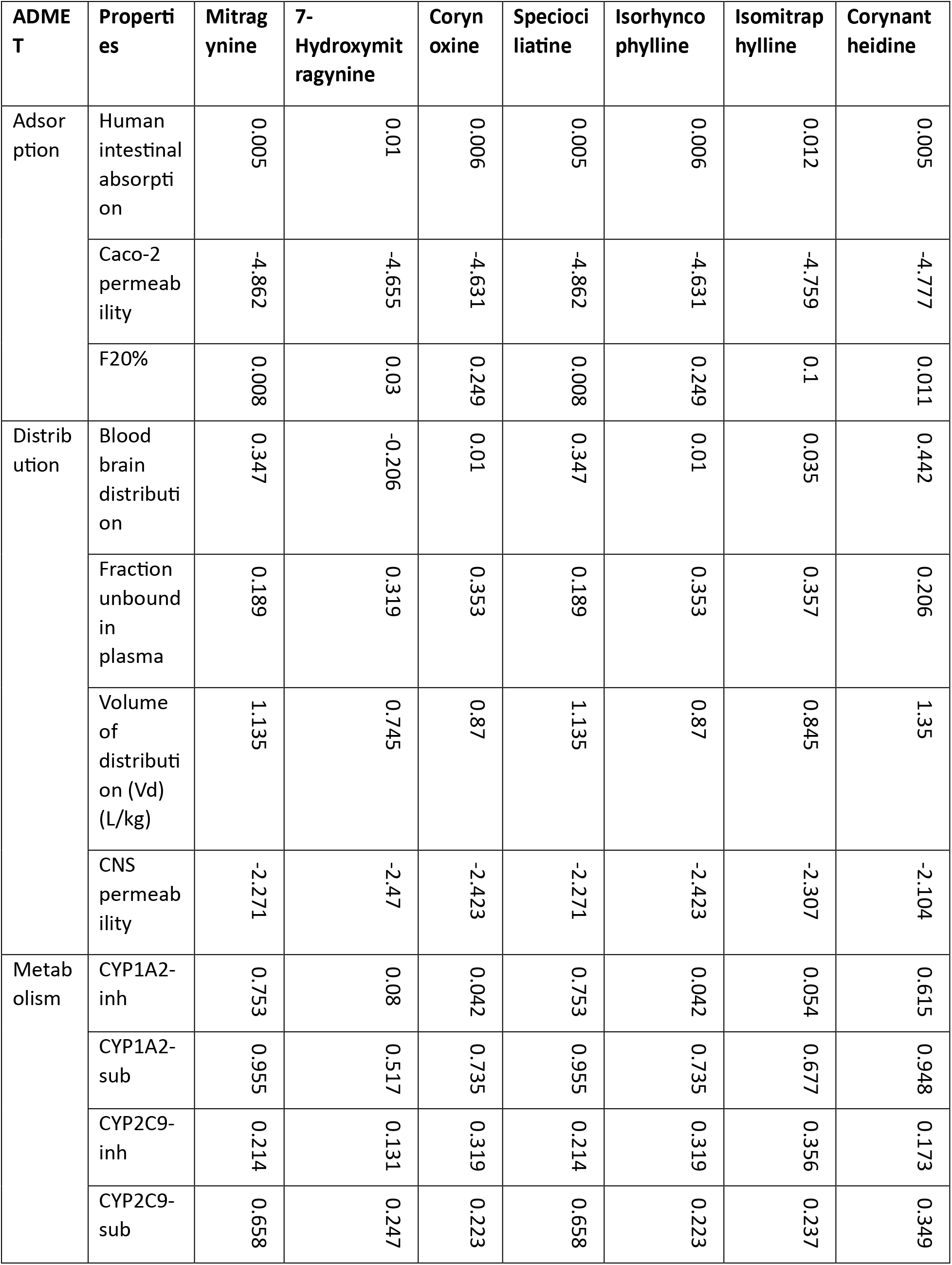
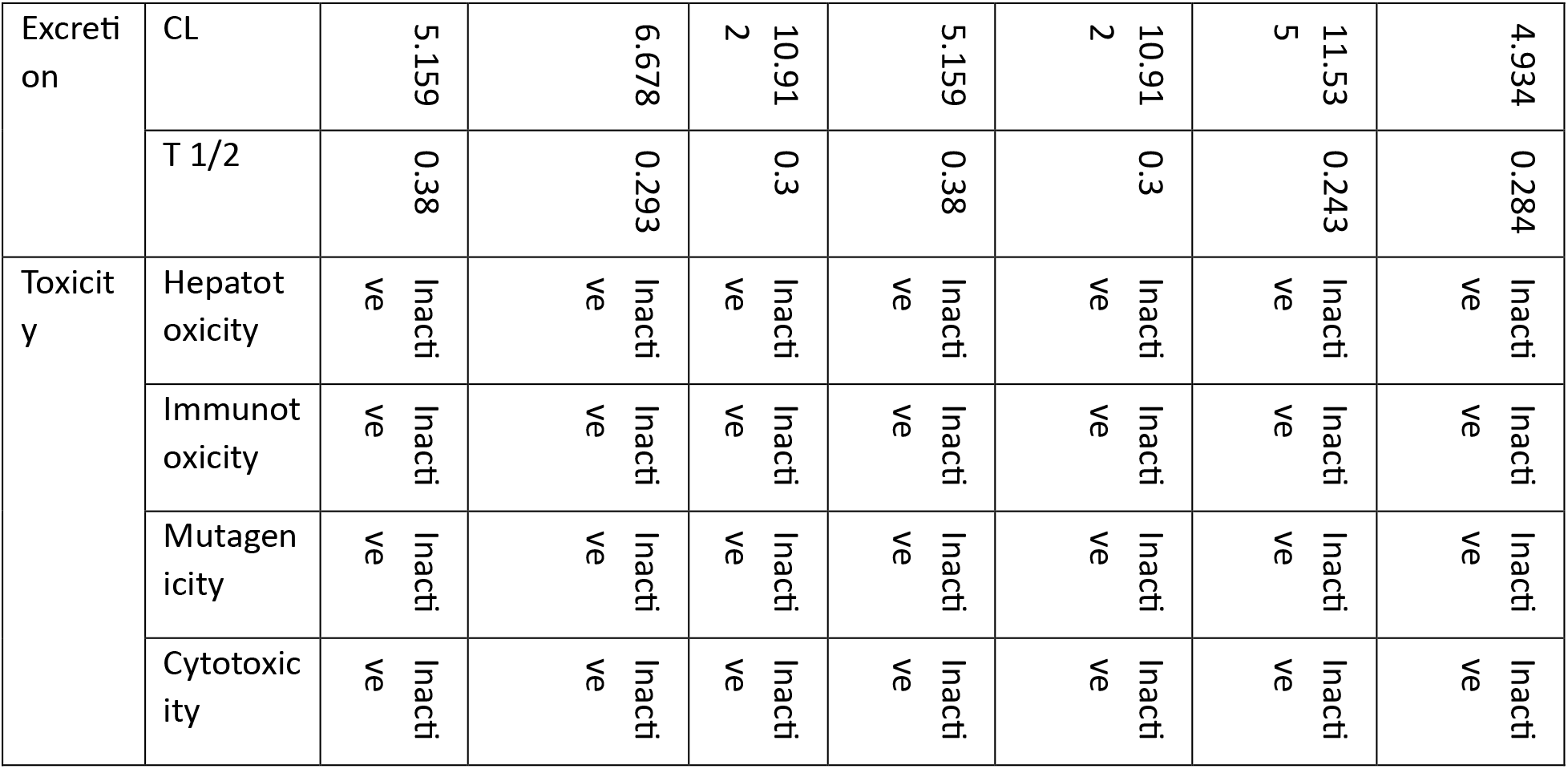
ADMET properties of seven compounds analyzed using ADMETlab 2.0, pkCSM and ProTox-II.

| ADMET | Properties | Mitragynine | 7-Hydroxymitragynine | Corynoxine | Speciociliatine | Isorhynchophylline | Isomitraphylline | Corynantheidine |
| --- | --- | --- | --- | --- | --- | --- | --- | --- |
| Adsorption | Human intestinal absorption | 0.005 | 0.01 | 0.006 | 0.005 | 0.006 | 0.012 | 0.005 |
|  | Caco-2 permeability | -4.862 | -4.655 | -4.631 | -4.862 | -4.631 | -4.759 | -4.777 |
|  | F20% | 0.008 | 0.03 | 0.249 | 0.008 | 0.249 | 0.1 | 0.011 |
| Distribution | Blood brain distribution | 0.347 | -0.206 | 0.01 | 0.347 | 0.01 | 0.035 | 0.442 |
|  | Fraction unbound in plasma | 0.189 | 0.319 | 0.353 | 0.189 | 0.353 | 0.357 | 0.206 |
|  | Volume of distribution (Vd) (L/kg) | 1.135 | 0.745 | 0.87 | 1.135 | 0.87 | 0.845 | 1.35 |
|  | CNS permeability | -2.271 | -2.47 | -2.423 | -2.271 | -2.423 | -2.307 | -2.104 |
| Metabolism | CYP1A2-inh | 0.753 | 0.08 | 0.042 | 0.753 | 0.042 | 0.054 | 0.615 |
|  | CYP1A2-sub | 0.955 | 0.517 | 0.735 | 0.955 | 0.735 | 0.677 | 0.948 |
|  | CYP2C9-inh | 0.214 | 0.131 | 0.319 | 0.214 | 0.319 | 0.356 | 0.173 |
|  | CYP2C9-sub | 0.658 | 0.247 | 0.223 | 0.658 | 0.223 | 0.237 | 0.349 |
| Excretion | CL | 5.159 | 6.678 | 10.912 | 5.159 | 10.912 | 11.535 | 4.934 |
|  | T 1/2 | 0.38 | 0.293 | 0.3 | 0.38 | 0.3 | 0.243 | 0.284 |
| Toxicity | Hepatotoxicity | Inactive | Inactive | Inactive | Inactive | Inactive | Inactive | Inactive |
|  | Immunotoxicity | Inactive | Inactive | Inactive | Inactive | Inactive | Inactive | Inactive |
|  | Mutagenicity | Inactive | Inactive | Inactive | Inactive | Inactive | Inactive | Inactive |
|  | Cytotoxicity | Inactive | Inactive | Inactive | Inactive | Inactive | Inactive | Inactive |

### 3.5 Active site and domain identification of DPP4

The active site of the selected DPP4 protein has an area of 4969.801 and contains 10120.087 amino acids (**Supplementary Figure 2**). Using the InterProScan web tool, two domains and an active residue of 605-635 position DPP4 are predicted [60]. The InterProScan predicted two DPP4 protein domains, the Dipeptidyl peptidase IV N-terminal region domain (108-478) and the Prolyl oligopeptidase family (559-763) (**Supplementary Figure 3**).

### 3.6 Molecular Docking and Interpretation of protein-ligands Interactions

Mitragynine, Corynantheidine, Corynoxine, and Speciociliatine (**Supplementary Figure 4**) from seven (7) compounds (**Supplementary Figure 5**) were selected for molecular docking using AutoDock vina to predict their binding affinity with the 5T4H_B receptor. The analysis predicted nine binding locations and the best one location was selected based on the lowest docking score and binding amino acid residues. Mitragynine, Corynantheidine, Corynoxine, and Speciociliatine showed docking energies as −7.5 kcal/mol, −7.7 kcal/mol, −7.1 kcal/mol, and −6.9 kcal/mol respectively. All the compounds were shown to be bound in the active cleft of the receptor (**Figure 5A**). In the case of Mitragynine one conventional hydrogen bond was predominantly formed with TYR B:752. The positions of TYR B:48, ARG B:125, LYS B:554, ASN B:562, TRP B:563, ALA B:564, TRP B:627, SER B:630, HIS B:740, GLY B:741, HIS B:748, acquired eleven van der Waals bonds, one carbon-hydrogen bond at ASP B:545, and one Pi-Pi Stacked and Pi-alkyl bond with TRP B:629 position (**Figure 5B**). For the compound Corynantheidine, there were observed two conventional hydrogen bonds with TYR B:48, TYR B:752, ten van der Waals interactions with ARG B:125, LYS B:554, ASN B:562, TRP B:563, ALA B:564, TRP B:627, SER B:630, HIS B:740, GLY B:741, HIS B:748, one carbon-hydrogen bond at ASP B:545, one Pi-Pi Stacked bond with TRP B:629, and one Pi-alkyl bond with TRP B:629 (**Figure 5B**). Corynoxine exhibited two conventional hydrogen bonds with VAL B:546, TRP B:629, seven van der Waals interactions with ASP B:545, TRP B:627, GLY B:628, GLY B:632, GLY B:741, HIS B:748, TYR B:752, one carbon-hydrogen bond at HIS B:740, one Pi-Pi Stacked bond with TYR B:547, one Pi-alkyl bond with RP B:629, and one Pi-cation bond with LYS B:554 (Figure 5B). Speciociliatine, the fourth molecule, had one conventional hydrogen bond at ARG B:125, six van der Waals interactions with PHE 273, ILE 347, HIS 363, ILE 411, PHE 413, one carbon-hydrogen bond at HIS B:740, four Pi-alkyl bonds with TYR B:631, TYR B:662, TYR B:666, HIS B:740, and one Pi-Pi Stacked at TYR B:547, one Pi-sigma bond with TRP B:629 position (**Figure 5B**). The interactions between the 5T4H_B protein and all four drug compounds shared the binding residues TRP B:629, HIS B:740, and GLY B:741.

**Figure 5:**
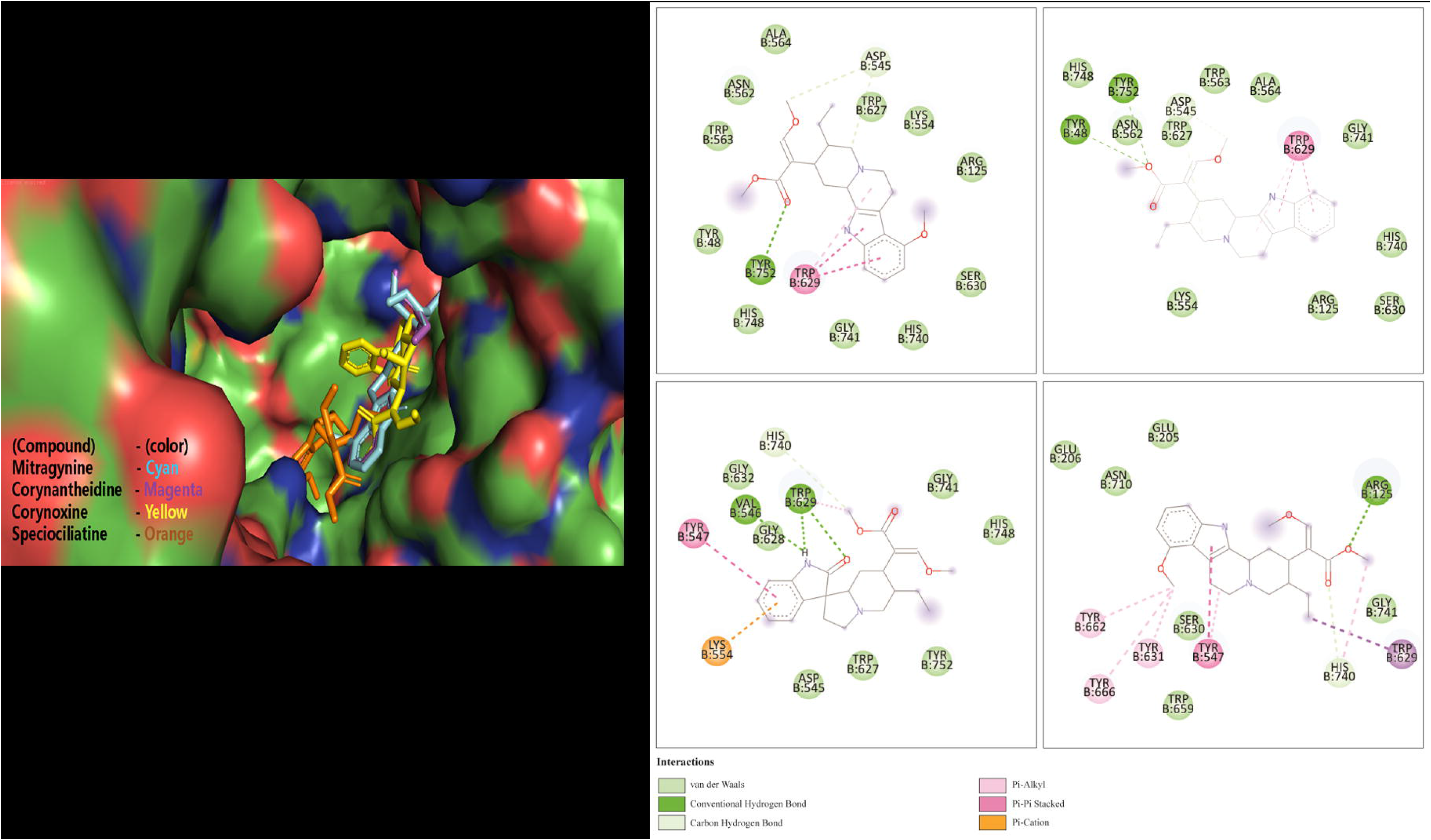
Drug binding to the receptor. 5A) Binding interactions of four compounds of Mitragyna speciosa with A chain of DPP4. The space filling model also showed the four compounds in the active site cleft of the B chain. Fig 5B) 2D interaction between the protein– ligand complex. Here, figure i) Mitragynine, ii)Corynantheidine, iii) Corynoxine, and iv) Speciociliatine showing the ligand contact with the protein DPP4 after molecular docking. Several types of interaction were shown by color with different amino acid residues.

### 3.7 Molecular Dynamics Simulation

The plot for the radius of gyration for the four (4) DPP4-drug complex Violet: DPP4-Corynantheidine complex; orange: DPP4-Corynoxine complex; green: DPP4-Mitragine complex; blue: DPP4-Speciociliatine complex demonstrates fluctuations over time **(Figure 6 A)**. The initial 30 nanoseconds are linear in trend except the DPP4-Corynantheidine complex which displays a slight upward change around 30 seconds. Afterward, all of the four (4) complexes displayed fluctuations over the next 40 seconds, among them DPP4-Corynantheidine complex and DPP4-Mitragine complex exhibited a higher rate of the radius of gyration from the beginning of the 30th second and maintained. Conversely, the final 30 seconds disregarded the linear trends with bias, the DPP4-Speciociliatine complex and DPP4-Mitragine complex were of high fluctuations and variations, whereas the DPP4-Corynoxine complex manifested almost a stable nature. Amidst them, the DPP4-Corynantheidine complex (violet) best maintained the sturdy and linear without much fluctuation or variation. The plot for the RMSD value for four (4) DPP4-drug complexes, RMSD plot showed that all DPP4-drug complexes gained stability after 15 ns, and became concurrent with the RMSD plot of DPPP4 for the next 25 ns **(Figure 6B)**. All of the complex was not found with any of the unusual fluctuations in the case of RMSF analysis except the DPP4-Mitragine complex which shows a gradual increase in the RMSD value with fluctuations. This shows that the three (3) complex was stable throughout the simulation run with minimal distortion. The fluctuation pattern of the DPP4-Corynantheidine complex, DPP4-Corynoxine complex, and DPP4-Speciociliatine complex was similar for the last 50 ns. The RMSF values for all four compounds are generally low, with most residues fluctuating between 0 and 0.4 nm **(Figure 6C)**. However, there are two regions of the protein that exhibit higher than average fluctuations. The first region is located between residue 250 and 200, where the RMSF value rises above 1 nm. This suggests that this region of the protein is more flexible or dynamic than other regions. The second region with higher than average fluctuation is located around residue 680, with an RMSF value below 0.5 nm but still higher than the average fluctuation. This region is not as flexible as the first one, but it still exhibits more motion than other parts of the protein. Overall, the RMSF graph provides a visual representation of how the flexibility or motion of each residue varies across the different compounds in the simulation. The higher RMSF values in certain regions of the protein suggest that these regions may be important for the function or stability of the protein, and further analysis may be necessary to understand their role. In the initial 40 ns, the DPP4-Corynantheidine complex shows a gradual increase in the SASA value with fluctuations, while another three DPP4-ligand complexes show a gradual decrease with fluctuation (**(Figure 6D)**. In the 85 ns time period, the DPP4-Corynantheidine complex showed the highest value of SASA and in the final 10 ns, all four (4) complexes displayed significant decreases in the SASA value with fluctuations.

**Figure 6:**
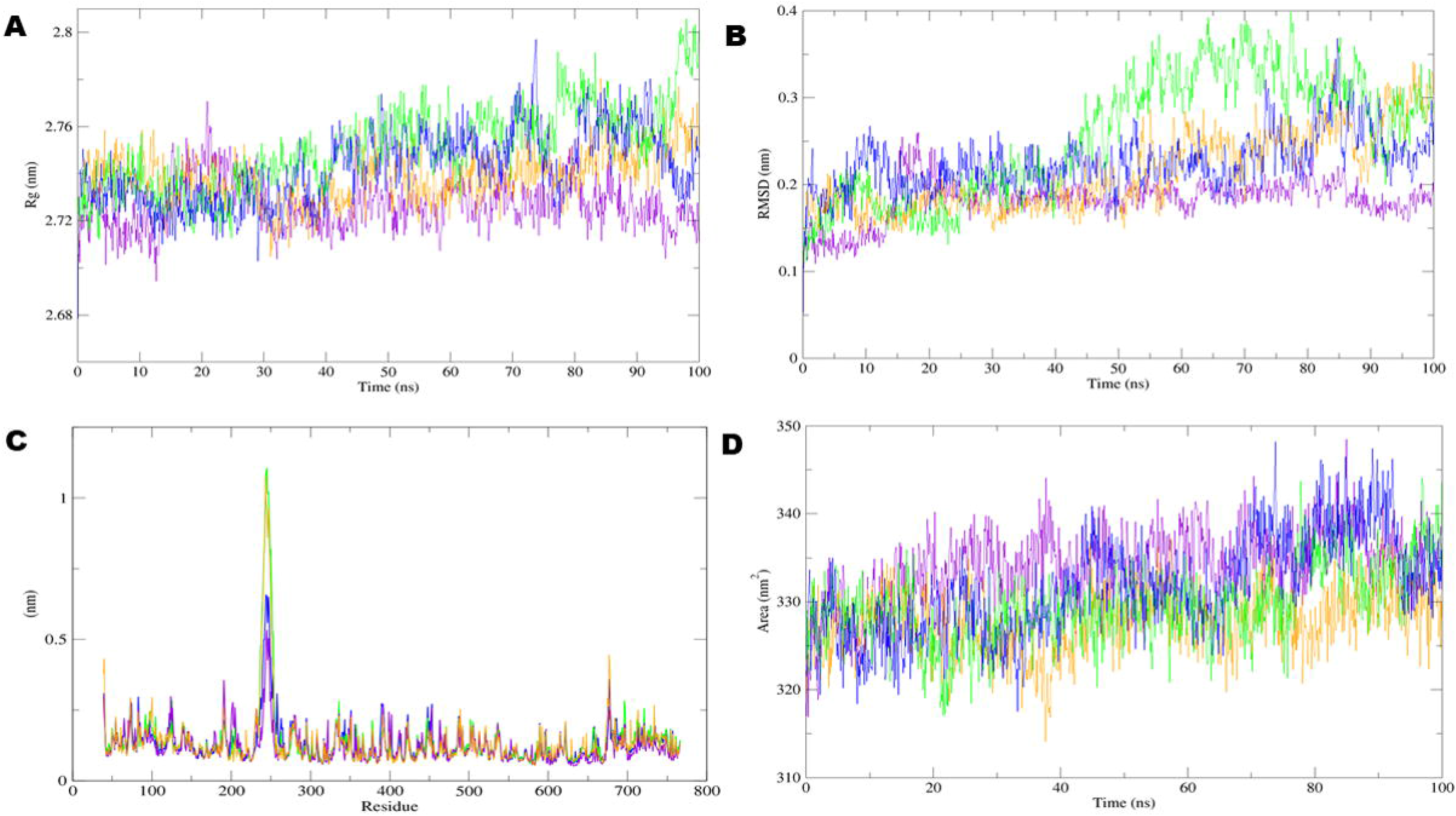
Molecular dynamics simulation of the DPP4 and ligand complex. A) Radius of gyration (Rg) for 100ns; B) 100ns-Root Mean Square Deviation (RMSD); C) Root-mean-square fluctuation (RMSF) for all residues; D) Solvent Accessible Surface Area (SASA), viz Violet: DPP4-Corynantheidine complex; orange: DPP4-Corynoxine complex; green: DPP4-Mitragine complex; blue: DPP4-Speciociliatine complex. Here, the Rg graph illustrates changes in protein compactness throughout the simulation, while RMSD indicates protein stability in the presence of drugs within the dynamic system. The RMSF graph portrays the mobility of the receptors when interacting with the drugs, and SASA reflects alterations in the solvent-accessible surface area under dynamic conditions.

## 4. Discussion

The study of compounds to facilitate drug design has received significant attention as a discipline of study in the past few decades. In recent years, computational compound screening has grown in popularity as an alternative method to the time-consuming and expensive screening process for investigating compounds [61], [62], [63]. Yet, no integrated approach has been performed on mouse models and computational screening of drug compounds. This integrated approach has therefore been validated experimentally with *M. speciosa*-induced diabetic mice (**Figure 1**). Consequently, *M. speciosa* (kratom) was applied to identify new bioactive drug candidates for an anti-diabetes study. For this reason, the alloxan drug was used in both male and female mice to cause the experimental diabetes. Alloxan is widely recognized to cause diabetes, which inhibits insulin secretion and induces the necrosis of β-cells [64]. After the induction of diabetes, the OGTT of both diabetic and non-diabetic mice was performed. The baseline of BGL non-diabetic mice showed non-significant changes in BGL after the 180 min treatment. However, the BGL of alloxan induced mice showed highly significant after the glucose level of 180 min (P>0.001) (**Figure 3A and 3B**). The area under the curve (AUC) also showed a significant difference in BGL (P<0.001) (**Figure 3C and 3D**). We then assessed the acute toxicity of *M. speciosa* methanolic extract. The mice had identical body weights, food consumption, and no behavioral changes, which suggests that *M. speciosa* extracts were safe for the dosages, but further research is required.

To examine how different extract doses affected the blood glucose levels in mice, both mice with and without diabetes were used in this study. The 200 mg/kg and 400 mg/kg extract doses were compared to glibenclamide, a commonly used anti-diabetic drug. When compared to the control in diabetic mice, the 200 mg/kg extract dose significantly decreased blood glucose levels (P < 0.001 in males and P < 0.001 in females). In the same manner, 400 mg/kg extract also reduced glucose levels significantly (P < 0.05 in males and P <0.0001 in females) compared to the control, but no significant effect was found between 200 mg/kgb.w and 400 mg/kgb.w of doses. In mice with or without diabetes, glibenclamide at 200 or 400 mg/kg did not significantly change the blood glucose levels (p > 0.05). As compared to the usual medication glibenclamide, the 200 mg/kg extract dose of *M. speciosa* demonstrated promising antidiabetic effects in diabetic mice (**Figure 4**). This happened because of the compounds in the extract that bind to cell receptors and release glucose for cellular use, lowering glucose levels in diabetic mice. After the evaluation of *M. speciosa* in diabetic mice as a potential antidiabetic plant, we retrieved 7 compounds from the *M. speciosa* plant using the PubChem database based on their reported therapeutic potential and underwent a series of computational analyses to evaluate their ADMET properties (**Table 1 and Supplementary Table 1**).

Using various experimental assays, the ADMET properties of the four compounds Mitragynine, Corynoxine, Speciociliatine, and Corynantheidine were evaluated and found to satisfy all parameters (**Table 1**). There is a high level of bioavailability shown by the F20% value, which ranges from 0.008 to 0.289. The value of intestinal absorption, which varies from 0.005 to 0.006, suggests that the body could absorb quite well. The distribution volume (Vd) values ranged from 0.87 to 1.35 L/kg, indicating that the compounds are well distributed throughout the body. However, the compounds have moderate to strong inhibitory effects on CYP enzymes, suggesting they may interact with other drugs metabolized by the same enzymes. Nonetheless, the compounds exhibited a low substrate potential for CYP enzymes, indicating that they are unlikely to be metabolized by these enzymes. The selected four compounds could easily be eliminated by the body after the administration, as indicated by CL values, ranging from 4.93 to 10.91. In addition, the compounds exhibited no toxicity in the hepatotoxicity, immunotoxicity, mutagenicity, and cytotoxicity assays, whereas only one compound was active in the carcinogenicity assay. Moreover, the selected four compounds showed strong binding energy ranging from −6.9 to −7.7 kcal/mol, having the interactions of van der Waals bonds, carbon-hydrogen bonds, and Pi-alkyl and Pi-Pi stacked bonds between the compounds and 5T4H_B protein. All four compounds shared the binding residues TRP B:629, HIS B:740, and GLY B:741 (**Supplementary Figure 1-5, Figure 5 and Table 2**).

**Table 2:** The properties of all four compounds.

| Properties | Mitragynine | Corynantheidine | Corynoxine | Speciociliatine |
| --- | --- | --- | --- | --- |
| **IUPAC Name** | methyl (E)-2-[(2S,3S,12bS)-3-ethyl-8-methoxy-1,2,3,4,6,7,12,12b-octahydroindolo[2,3-a]quinolizin-2-yl]-3-methoxyprop-2-enoate | methyl (E)-2-[(2S,3S,12bS)-3-ethyl-1,2,3,4,6,7,12,12b-octahydroindolo[2,3-a]quinolizin-2-yl]-3-methoxyprop-2-enoate | methyl (E)-2-[(3S,6'S,7'S,8'aS)-6'-ethyl-2-oxospiro[1H-indole-3,1'-3,5,6,7,8,8a-hexahydro-2H-indolizine]-7'-yl]-3-methoxyprop-2-enoate | methyl (E)-2-[(2S,3S,12bR)-3-ethyl-8-methoxy-1,2,3,4,6,7,12,12b-octahydroindolo[2,3-a]quinolizin-2-yl]-3-methoxyprop-2-enoate |
| **Molecular Weight** | 398.5 | 368.5 | 384.5 | 398.5 |
| **Hydrogen Bond Donor** | 1 | 1 | 1 | 1 |
| **Hydrogen Bond Acceptor** | 5 | 4 | 5 | 5 |
| **Docking Energy** | -7.5 kcal/mol | -7.7 kcal/mol | -7.1 kcal/mol | -6.9 kcal/mol |
| **Solubility (log mol/L)** | -4.181 | -4.115 | -3.709 | -4.181 |
| **Binding residues of DPP4 (B chain)** | TYR B:48, ARG B:125, ASP B:545, LYS B:554, ASN B:562, TRP B:563, ALA B:564, TRP B:627, TRP B:629, SER B:630, HIS B:740, GLY B:741, HIS B:748, TYR B:752 | TYR B:48, ARG B:125, ASP B:545, LYS B:554, ASN B:562, TRP B:563, ALA B:564, TRP B:627, TRP B:629, SER B:630, HIS B:740, GLY B:741, HIS B:748, TYR B:752 | ASP B:545, VAL B:546, TYR B:547, LYS B:554, TRP B:627, GLY B:628, TRP B:629, GLY B:632, HIS B:740, GLY B:741, HIS B:748, TYR B:752 | ARG B:125, GLU B:205, GLU B:206, TYR B:547, TRP B:629, SER B:630, TYR B:631, TRP B:659, TYR B:662, TYR B:666 ASN B:710, HIS B:740, GLY B:741 |

Furthermore, the values of molecular dynamics such as Rg, RMSD, RMSF, and SASA have shown fluctuations at different time points at 100ns. Several Molecular dynamics simulation studies have been done previously to investigate the stability of molecules over time [65], [66], [67], [68]. However, these values remained constant throughout time, demonstrating that the compound was firmly bound to the receptor (**Figure 6**). Consequently, upon experimental confirmation, it may be suggested that Mitragynine and Corynantheidine could potentially serve as oral medications for DM based on their binding residues at the active site, minimal toxicity, excellent docking scores, and low molecular weight. The virtual screening of natural plant compounds from *M. speciosa* (Mitragynine, Corynantheidine, Corynoxine, and Speciociliatine) revealed their potential as oral medications based on *in silico* analysis. The antidiabetic properties of the plant extract were confirmed in diabetic animal models. To sum up, our study recommends four potential antidiabetic compounds against diabetes (**Figure 7**).

**Figure 7:**
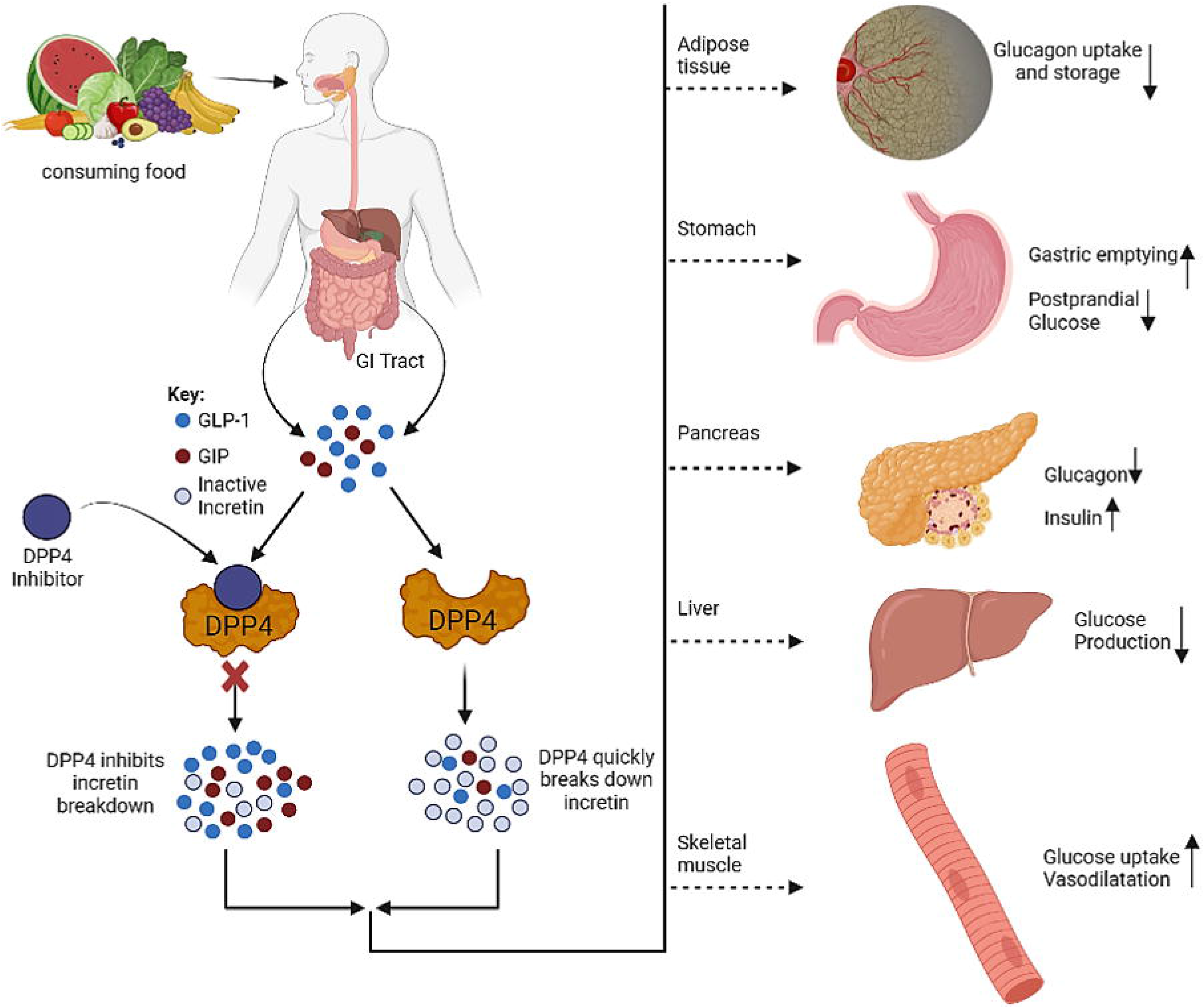
The effect of DPP4 inhibitor and its potential effects in diabetes. Dipeptidyl-peptidase-4 (DPP-4) inhibitors exert their anti-hyperglycemic effects by preventing the deactivation of incretin hormones, such as glucagon-like peptide-1 (GLP-1) and glucose-dependent insulinotropic polypeptide (GIP). These inhibitors impact glucose regulation through various mechanisms in adipose tissues, stomach, pancreas, liver, and skeletal muscles respectively.

## 5. Limitations of the study

Although the *in silico* and *in vivo* yielded promising findings, the present study has several shortcomings. Firstly, no compounds were extracted and identified using chromatographic techniques. Secondly, there is a possibility that the results from *in silico* experiments may not accurately reflect the actual behavior of the compounds’ clinical trials. Lastly, the study only evaluated the potential therapeutic effects of the *M. speciosa* plant extracts on diabetes and did not investigate their effects on other diseases.

## 6. Conclusion

The study confirmed that the methanolic extract of *M. speciosa* and the four potential compounds have insulin mimicking activity. Further investigation is required to ascertain the optimal dosage and the mode of action. These results imply that plant extracts from *M. speciosa* and their constituents, including speciociliatine, mitragynine, corynantheidine, and corynoxine, may offer a novel approach to the treatment of diabetes.

## 8. Author Contribution

**Mohammad Uzzal Hossain, Md. Shahadat Hossain, Shajib Dey, A.B.Z Naimur Rahman**, **Zeshan Mahmud Chowdhury**: Conceptualization; data retrieval; sequence analysis; laboratory experiments; writing – original draft; **Arittra Bhattacharjee**: writing – review and editing; **Ishtiaque Ahammad**: manuscript preparations; **Md. Nazrul Islam**: data curation; methodology; **Md. Moniruzzaman:** sample and data collection; **Md. Billal Hosen:** data curation; methodology; **Istiaq Ahmed**: data curation; methodology; **Keshob Chandra Das:** supervision; validation; **Chaman Ara Keya:** supervision; validation; **Md. Salimullah:** project administration; validation; supervision.

## Supporting information

https://drive.google.com/drive/folders/1Kq8P17uJVV2Q3qktVbPGOVDFsHJ2rcXy?usp=sharing

## Reference

[1] A. Ramachandran, “Know the signs and symptoms of diabetes.,” Indian J Med Res, vol. 140, no. 5, pp. 579–81, Nov. 2014.

[2] J. M. Forbes and M. E. Cooper, “Mechanisms of Diabetic Complications,” Physiol Rev, vol. 93, no. 1, pp. 137–188, Jan. 2013, doi: 10.1152/physrev.00045.2011.

[3] O. Ali, “Genetics of type 2 diabetes,” World J Diabetes, vol. 4, no. 4, p. 114, 2013, doi: 10.4239/wjd.v4.i4.114.

[4] A. D. Association, “2. Classification and Diagnosis of Diabetes: Standards of Medical Care in Diabetes—2021,” Diabetes Care, vol. 44, no. Supplement_1, pp. S15–S33, Jan. 2021, doi: 10.2337/DC21-S002.

[5] B. Silver et al., “EADSG Guidelines: Insulin Therapy in Diabetes.,” Diabetes Ther, vol. 9, no. 2, pp. 449–492, Apr. 2018, doi: 10.1007/s13300-018-0384-6.

[6] P. E. Cryer, S. N. Davis, and H. Shamoon, “Hypoglycemia in Diabetes,” Diabetes Care, vol. 26, no. 6, pp. 1902–1912, Jun. 2003, doi: 10.2337/diacare.26.6.1902.

[7] J. Rosenstock, S. L. Schwartz, C. M. Clark, G. D. Park, D. W. Donley, and M. B. Edwards, “Basal Insulin Therapy in Type 2 Diabetes28-week comparison of insulin glargine (HOE 901) and NPH insulin,” Diabetes Care, vol. 24, no. 4, pp. 631–636, Apr. 2001, doi: 10.2337/DIACARE.24.4.631.

[8] C. Sorli and M. K. Heile, “Identifying and meeting the challenges of insulin therapy in type 2 diabetes,” J Multidiscip Healthc, p. 267, Jul. 2014, doi: 10.2147/JMDH.S64084.

[9] A. Chaudhury et al., “Clinical Review of Antidiabetic Drugs: Implications for Type 2 Diabetes Mellitus Management,” Front Endocrinol (Lausanne), vol. 8, Jan. 2017, doi: 10.3389/fendo.2017.00006.

[10] G. Roglic, “WHO Global report on diabetes: A summary,” Int J Noncommun Dis, vol. 1, no. 1, p. 3, 2016, doi: 10.4103/2468-8827.184853.

[11] J. da Rocha Fernandes et al., “IDF Diabetes Atlas estimates of 2014 global health expenditures on diabetes,” Diabetes Res Clin Pract, vol. 117, pp. 48–54, Jul. 2016, doi: 10.1016/j.diabres.2016.04.016.

[12] M. Ekor, “The growing use of herbal medicines: issues relating to adverse reactions and challenges in monitoring safety,” Front Pharmacol, vol. 4, 2014, doi: 10.3389/fphar.2013.00177.

[13] H. Choudhury et al., “An update on natural compounds in the remedy of diabetes mellitus: A systematic review,” J Tradit Complement Med, vol. 8, no. 3, pp. 361–376, Jul. 2018, doi: 10.1016/j.jtcme.2017.08.012.

[14] M. L. Willcox, C. Elugbaju, M. Al-Anbaki, M. Lown, and B. Graz, “Effectiveness of Medicinal Plants for Glycaemic Control in Type 2 Diabetes: An Overview of Meta-Analyses of Clinical Trials,” Front Pharmacol, vol. 12, 2021, doi: 10.3389/fphar.2021.777561.

[15] B. Dineshkumar B., M. Analava M., and M. Manjunatha M., “Antidiabetic and hypolipidaemic effects of few common plants extract in Type 2 diabetic patients at Bengal,” International Journal of Diabetes and Metabolism, vol. 18, no. 2, pp. 59–65, 2010, doi: 10.1159/000497694.

[16] G. Bjørklund, M. Dadar, L. Pivina, M. D. Doşa, Y. Semenova, and J. Aaseth, “The Role of Zinc and Copper in Insulin Resistance and Diabetes Mellitus,” Curr Med Chem, vol. 27, no. 39, pp. 6643–6657, Nov. 2020, doi: 10.2174/0929867326666190902122155.

[17] A. Al-Brakati, “Protective Effect of Garlic against Diabetic Retinopathy in Adult Albino Rats,” 2016. [Online]. Available: https://api.semanticscholar.org/CorpusID:35413704

[18] G. Velu, V. Palanichamy, and A. P. Rajan, “Phytochemical and Pharmacological Importance of Plant Secondary Metabolites in Modern Medicine,” in *Bioorganic Phase in Natural Food: An Overview*, Cham: Springer International Publishing, 2018, pp. 135–156. doi: 10.1007/978-3-319-74210-6_8.

[19] Y. H. Gonfa, F. Beshah, M. G. Tadesse, A. Bachheti, and R. K. Bachheti, “Phytochemical investigation and potential pharmacologically active compounds of Rumex nepalensis: an appraisal,” Beni Suef Univ J Basic Appl Sci, vol. 10, no. 1, p. 18, Dec. 2021, doi: 10.1186/s43088-021-00110-1.

[20] J. R. Shaikh and M. Patil, “Qualitative tests for preliminary phytochemical screening: An overview,” Int J Chem Stud, vol. 8, no. 2, pp. 603–608, Mar. 2020, doi: 10.22271/chemi.2020.v8.i2i.8834.

[21] Y. Gavamukulya, F. Abou-Elella, F. Wamunyokoli, and H. AEl-Shemy, “Phytochemical screening, anti-oxidant activity and in vitro anticancer potential of ethanolic and water leaves extracts of Annona muricata (Graviola),” Asian Pac J Trop Med, vol. 7, pp. S355–S363, Sep. 2014, doi: 10.1016/S1995-7645(14)60258-3.

[22] K. Prabu, A. Rajasekaran, D. Bharathi, and S. Ramalakshmi, “Anti-oxidant activity, phytochemical screening and HPLC profile of rare endemic Cordia diffusa,” J King Saud Univ Sci, vol. 31, no. 4, pp. 724–727, Oct. 2019, doi: 10.1016/j.jksus.2018.04.025.

[23] Bindu Jacob and Narendhirakannan R.T., “Role of medicinal plants in the management of diabetes mellitus: a review,” 3 Biotech, vol. 9, no. 1, p. 4, Jan. 2019, doi: 10.1007/s13205-018-1528-0.

[24] N. Sharma and K. S. Bora, “Role of Medicinal Plants in the Management of Diabetes Mellitus: A Review,” J Pharm Res Int, pp. 2196–2207, Dec. 2021, doi: 10.9734/jpri/2021/v33i60B34864.

[25] W. C. Prozialeck, J. K. Jivan, and S. V Andurkar, “Pharmacology of kratom: an emerging botanical agent with stimulant, analgesic and opioid-like effects.,” J Am Osteopath Assoc, vol. 112, no. 12, pp. 792–9, Dec. 2012.

[26] E. Cinosi et al., “Following ‘the Roots’ of Kratom (Mitragyna speciosa): The Evolution of an Enhancer from a Traditional Use to Increase Work and Productivity in Southeast Asia to a Recreational Psychoactive Drug in Western Countries.,” Biomed Res Int, vol. 2015, p. 968786, 2015, doi: 10.1155/2015/968786.

[27] K. G. M. M. Alberti et al., “Harmonizing the metabolic syndrome: a joint interim statement of the International Diabetes Federation Task Force on Epidemiology and Prevention; National Heart, Lung, and Blood Institute; American Heart Association; World Heart Federation; International Atherosclerosis Society; and International Association for the Study of Obesity.,” Circulation, vol. 120, no. 16, pp. 1640–5, Oct. 2009, doi: 10.1161/CIRCULATIONAHA.109.192644.

[28] L. Chen, S. Fei, and O. J. Olatunji, “LC/ESI/TOF-MS Characterization, Anxiolytic and Antidepressant-like Effects of Mitragyna speciosa Korth Extract in Diabetic Rats,” Molecules, vol. 27, no. 7, p. 2208, Mar. 2022, doi: 10.3390/molecules27072208.

[29] Y. S. Goh, T. Karunakaran, V. Murugaiyah, R. Santhanam, M. H. Abu Bakar, and S. Ramanathan, “Accelerated Solvent Extractions (ASE) of Mitragyna speciosa Korth. (Kratom) Leaves: Evaluation of Its Cytotoxicity and Antinociceptive Activity,” Molecules, vol. 26, no. 12, p. 3704, Jun. 2021, doi: 10.3390/molecules26123704.

[30] J. Ganugapati and S. Swarna, “MOLECULAR DOCKING STUDIES OF ANTIDIABETIC ACTIVITY OF CINNAMON COMPOUNDS,” Asian Journal of Pharmaceutical and Clinical Research, vol. 7, pp. 31–34, 2014, [Online]. Available: https://api.semanticscholar.org/CorpusID:83098482

[31] D. Ahmed, V. Kumar, M. Sharma, and A. Verma, “Target guided isolation, in-vitro antidiabetic, antioxidant activity and molecular docking studies of some flavonoids from Albizzia Lebbeck Benth. bark,” BMC Complement Altern Med, vol. 14, no. 1, p. 155, 2014, doi: 10.1186/1472-6882-14-155.

[32] S. C. Eastlack, E. M. Cornett, and A. D. Kaye, “Kratom-Pharmacology, Clinical Implications, and Outlook: A Comprehensive Review.,” Pain Ther, vol. 9, no. 1, pp. 55–69, Jun. 2020, doi: 10.1007/s40122-020-00151-x.

[33] J. E. Adkins, E. W. Boyer, and C. R. McCurdy, “Mitragyna speciosa, a psychoactive tree from Southeast Asia with opioid activity,” Curr Top Med Chem, vol. 11, no. 9, p. 1165–1175, 2011, doi: 10.2174/156802611795371305.

[34] P. Zhang, W. Wei, X. Zhang, C. Wen, C. Ovatlarnporn, and O. J. Olatunji, “Antidiabetic and antioxidant activities of Mitragyna speciosa (kratom) leaf extract in type 2 diabetic rats,” Biomedicine & Pharmacotherapy, vol. 162, p. 114689, 2023, doi: 10.1016/j.biopha.2023.114689.

[35] G. P. S. Kumar, P. Arulselvan, D. S. Kumar, and S. P. Subramanian, “Anti-Diabetic Activity of Fruits of Terminalia chebula on Streptozotocin Induced Diabetic Rats,” Journal of Health Science, vol. 52, no. 3, pp. 283–291, 2006, doi: 10.1248/jhs.52.283.

[36] T. Miura, T. Koike, and T. Ishida, “Antidiabetic Activity of Green Tea (Thea sinensis L.) in Genetically Type 2 Diabetic Mice,” Journal of Health Science, vol. 51, no. 6, pp. 708–710, 2005, doi: 10.1248/jhs.51.708.

[37] A. N. Nagappa, P. A. Thakurdesai, N. Venkat Rao, and J. Singh, “Antidiabetic activity of Terminalia catappa Linn fruits,” J Ethnopharmacol, vol. 88, no. 1, pp. 45–50, Sep. 2003, doi: 10.1016/S0378-8741(03)00208-3.

[38] National Research Council (U.S.). Committee for the Update of the Guide for the Care and Use of Laboratory Animals. and Institute for Laboratory Animal Research (U.S.), Guide for the care and use of laboratory animals. National Academies Press, 2011.

[39] B. Ragavan and S. Krishnakumari, “Antidiabetic effect of T. arjuna bark extract in alloxan induced diabetic rats,” Indian Journal of Clinical Biochemistry, vol. 21, no. 2, pp. 123–128, Sep. 2006, doi: 10.1007/BF02912926.

[40] Oecd, “OECD GUIDELINE FOR TESTING OF CHEMICALS Acute Oral Toxicity-Up-and-Down Procedure INTRODUCTION,” 2001.

[41] C. Bürger, D. R. Fischer, D. A. Cordenunzzi, A. P. de B. Batschauer, V. Cechinel Filho, and A. R. dos S. Soares, “Acute and subacute toxicity of the hydroalcoholic extract from Wedelia paludosa (Acmela brasiliensis) (Asteraceae) in mice.,” J Pharm Pharm Sci, vol. 8, no. 2, pp. 370–3, Aug. 2005.

[42] E. N. de Carvalho, N. A. S. de Carvalho, and L. M. Ferreira, “Experimental model of induction of diabetes mellitus in rats,” Acta Cir Bras, vol. 18, no. spe, pp. 60–64, 2003, doi: 10.1590/S0102-86502003001100009.

[43] N. Kamalakkannan and P. S. M. Prince, “Hypoglycaemic effect of water extracts of Aegle marmelos fruits in streptozotocin diabetic rats,” J Ethnopharmacol, vol. 87, no. 2–3, pp. 207–210, Aug. 2003, doi: 10.1016/S0378-8741(03)00148-X.

[44] A. Gidado, D. A. Ameh, and S. E. Atawodi, “Effect of Nauclea latifolia leaves aqueous extracts on blood glucose levels of normal and alloxan-induced diabetic rats,” Afr J Biotechnol, vol. 4, no. 1, pp. 91–93, 2005.

[45] L. Pari and S. Venkateswaran, “Effect of an aqueous extract of Phaseolus vulgaris on the properties of tail tendon collagen of rats with streptozotocin-induced diabetes,” Brazilian Journal of Medical and Biological Research, vol. 36, no. 7, pp. 861–870, Jul. 2003, doi: 10.1590/S0100-879X2003000700006.

[46] S. Kim et al., “PubChem Substance and Compound databases,” Nucleic Acids Res, vol. 44, no. D1, pp. D1202–D1213, Jan. 2016, doi: 10.1093/nar/gkv951.

[47] H. M. Berman, “The Protein Data Bank,” Nucleic Acids Res, vol. 28, no. 1, pp. 235–242, Jan. 2000, doi: 10.1093/nar/28.1.235.

[48] G. Xiong et al., “ADMETlab 2.0: an integrated online platform for accurate and comprehensive predictions of ADMET properties,” Nucleic Acids Res, vol. 49, no. W1, pp. W5–W14, Jul. 2021, doi: 10.1093/nar/gkab255.

[49] D. E. V. Pires, T. L. Blundell, and D. B. Ascher, “pkCSM: Predicting Small-Molecule Pharmacokinetic and Toxicity Properties Using Graph-Based Signatures,” J Med Chem, vol. 58, no. 9, pp. 4066–4072, May 2015, doi: 10.1021/acs.jmedchem.5b00104.

[50] P. Banerjee, A. O. Eckert, A. K. Schrey, and R. Preissner, “ProTox-II: a webserver for the prediction of toxicity of chemicals,” Nucleic Acids Res, vol. 46, no. W1, pp. W257–W263, Jul. 2018, doi: 10.1093/nar/gky318.

[51] N. Guex and M. C. Peitsch, “SWISS-MODEL and the Swiss-Pdb Viewer: An environment for comparative protein modeling,” Electrophoresis, vol. 18, no. 15, pp. 2714–2723, 1997, doi: 10.1002/elps.1150181505.

[52] M. D. Hanwell, D. E. Curtis, D. C. Lonie, T. Vandermeersch, E. Zurek, and G. R. Hutchison, “Avogadro: an advanced semantic chemical editor, visualization, and analysis platform,” J Cheminform, vol. 4, no. 1, p. 17, Dec. 2012, doi: 10.1186/1758-2946-4-17.

[53] T. A. Binkowski, “CASTp: Computed Atlas of Surface Topography of proteins,” Nucleic Acids Res, vol. 31, no. 13, pp. 3352–3355, Jul. 2003, doi: 10.1093/nar/gkg512.

[54] B. J. McConkey, V. Sobolev, and M. Edelman, “The performance of current methods in ligand– protein docking,” Curr Sci, vol. 83, no. 7, pp. 845–856, 2002, [Online]. Available: http://www.jstor.org/stable/24107087

[55] J. Eberhardt, D. Santos-Martins, A. F. Tillack, and S. Forli, “AutoDock Vina 1.2.0: New Docking Methods, Expanded Force Field, and Python Bindings,” J Chem Inf Model, vol. 61, no. 8, pp. 3891–3898, Aug. 2021, doi: 10.1021/acs.jcim.1c00203.

[56] L. L. C. Schrödinger and W. DeLano, “PyMOL.” [Online]. Available: http://www.pymol.org/pymol

[57] D. Systèmes, “‘BIOVIA Discovery Studio’ ‘Dassault Syst mes BIOVIA, Discovery Studio Modeling Environment, Release 2017’ Dassault Syst mes,” 2016.

[58] L. D. Schuler, X. Daura, and W. F. van Gunsteren, “An improved GROMOS96 force field for aliphatic hydrocarbons in the condensed phase,” J Comput Chem, vol. 22, no. 11, pp. 1205–1218, Aug. 2001, doi: 10.1002/jcc.1078.

[59] IBM Corp, “IBM SPSS Statistics for Windows.” IBM Corp, Armonk, NY, 2020.

[60] E. Quevillon et al., “InterProScan: protein domains identifier.,” Nucleic Acids Res, vol. 33, no. Web Server issue, pp. W116–20, Jul. 2005, doi: 10.1093/nar/gki442.

[61] J. Bajorath, “Integration of virtual and high-throughput screening,” Nat Rev Drug Discov, vol. 1, no. 11, pp. 882–894, Nov. 2002, doi: 10.1038/nrd941.

[62] A. Chien, I. Foster, and D. Goddette, “Grid technologies empowering drug discovery.,” Drug Discov Today, vol. 7, no. 20 Suppl, pp. S176–80, Oct. 2002, doi: 10.1016/s1359-6446(02)02369-3.

[63] M. U. Hossain et al., “Treating Diabetes Mellitus: Pharmacophore Based Designing of Potential Drugs fromGymnema sylvestreagainst Insulin Receptor Protein,” BioMed Research International, vol. 2016, pp. 1–14, Jan. 2016, doi: 10.1155/2016/3187647.

[64] O. M. Ighodaro, A. M. Adeosun, and O. Akinloye, “Alloxan-induced diabetes, a common model for evaluating the glycemic-control potential of therapeutic compounds and plants extracts in experimental studies,” Medicina-lithuania, vol. 53, no. 6, pp. 365–374, Jan. 2017, doi: 10.1016/j.medici.2018.02.001.

[65] M. U. Hossain et al., “Novel mutations in NSP-1 and PLPro of SARS-CoV-2 NIB-1 genome mount for effective therapeutics,” Journal of Genetic Engineering and Biotechnology, vol. 19, no. 1, Apr. 2021, doi: 10.1186/s43141-021-00152-z.

[66] Z. M. Chowdhury et al., “Exploration of Streptococcus core genome to reveal druggable targets and novel therapeutics against S. pneumoniae,” PLOS ONE, vol. 17, no. 8, p. e0272945, Aug. 2022, doi: 10.1371/journal.pone.0272945.

[67] I. Ahammad et al., “Impact of highly deleterious non-synonymous polymorphisms on GRIN2A protein’s structure and function,” PLOS ONE, vol. 18, no. 6, p. e0286917, Jun. 2023, doi: 10.1371/journal.pone.0286917.

[68] A. Al-Mamun, et al., “Pharmacoinformatics and molecular dynamics simulation approach to identify anti-diarrheal potentials of Centella asiatica (L.) Urb. against Vibrio cholerae,” Journal of Biomolecular Structure & Dynamics, pp. 1–14, Mar. 2023, doi: 10.1080/07391102.2023.2191736.

