## Supplementary figures and images for "*In vivo and in silico* experiment on *Mitragyna speciosa* offers new insight of antidiabetic potentials"

### Supplementary figure 1.tif

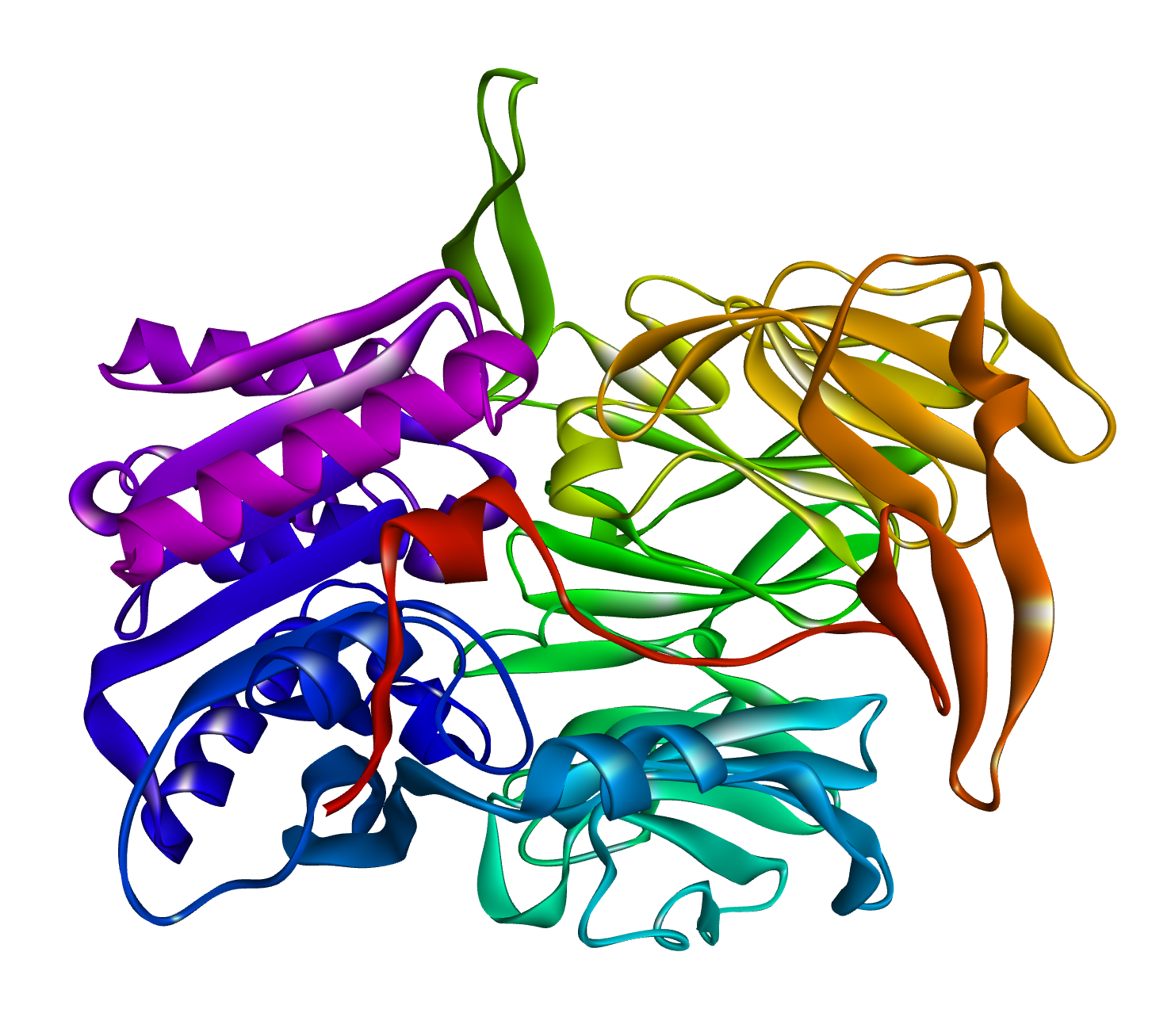

### Supplementary Figure 2.tif

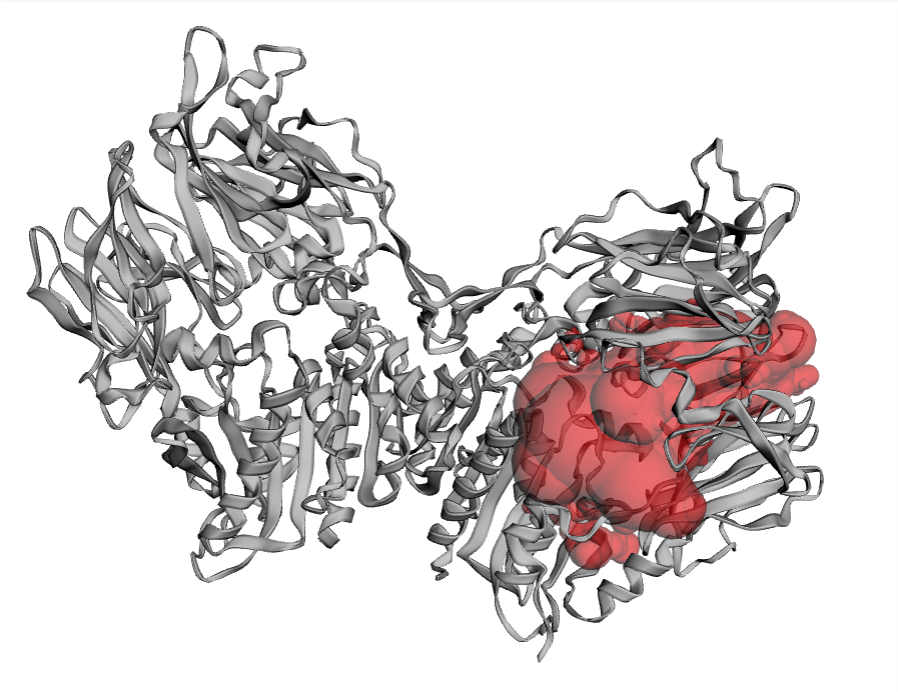

### Supplementary Figure 3.png

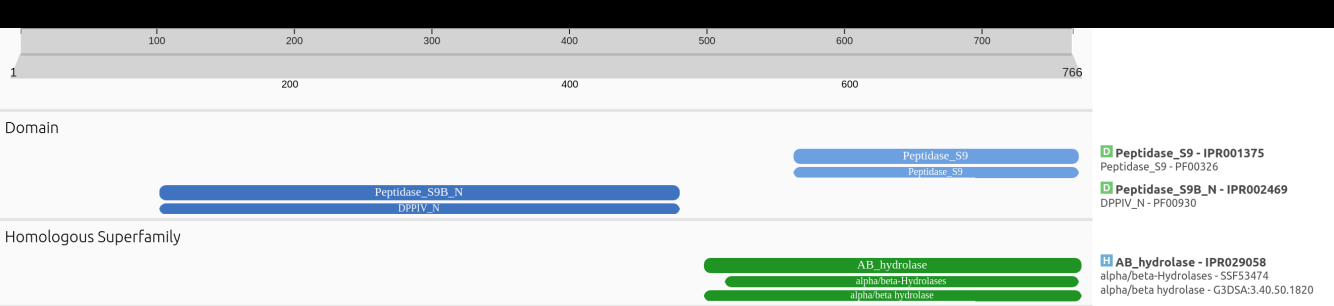

### Supplementary figure 4.tif

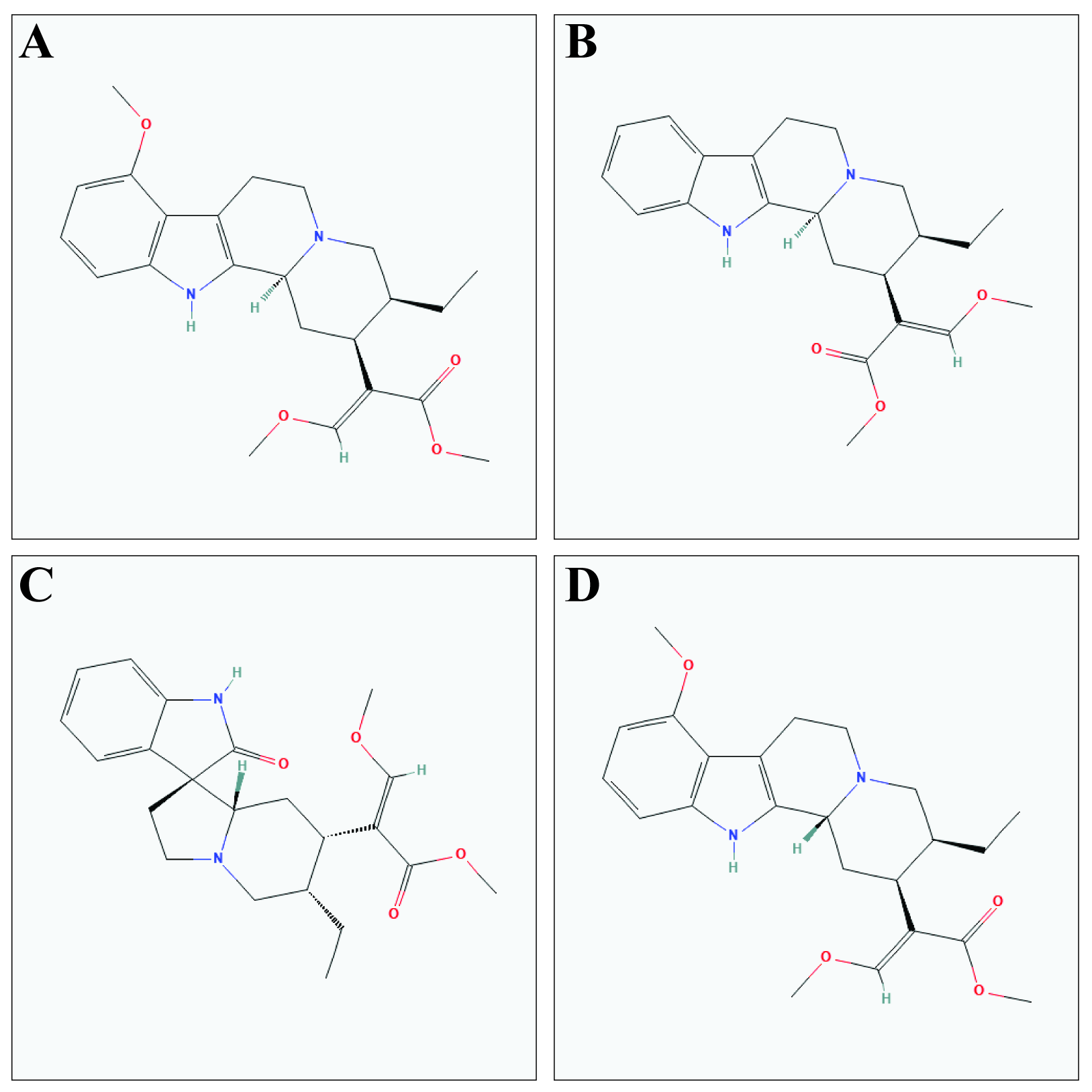

### Supplementary figure 5.tif

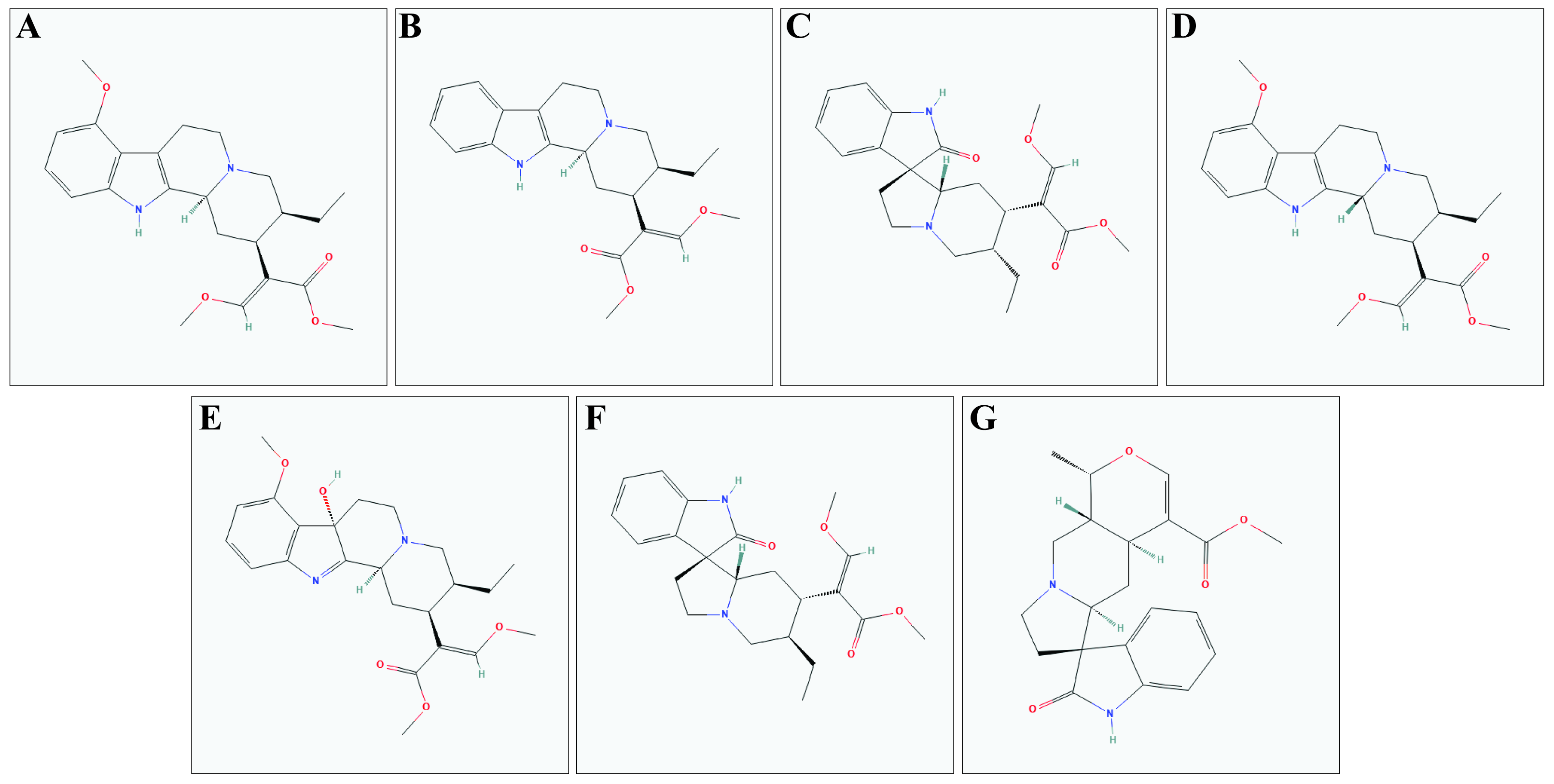
